# A conformational switch driven by phosphorylation regulates Ykt6 activity in macroautophagy

**DOI:** 10.1101/2020.03.15.992727

**Authors:** Kaitlyn McGrath, Mykola Dergai, Shivani Agarwal, Daayun Chung, Damian B. van Rossum, Aishwarya Shevade, Sergei Kuchin, Sofia Zaichick, Jeffrey N Savas, Dirk Fasshauer, Gabriela Caraveo

**Affiliations:** Department of Neurology, Feinberg School of Medicine, Northwestern University, Chicago, IL, USA; Department of Fundamental Neurosciences, University of Lausanne, Lausanne, Switzerland; Division of Experimental Pathology, Department of Pathology, Penn State College of Medicine, Hershey, PA, USA; The Jake Gittlen Laboratories for Cancer Research, Penn State College of Medicine, Hershey, PA; Department of Biological Sciences, University of Wisconsin Milwaukee, Milwaukee, WI, USA

## Abstract

Membrane fusion, an essential process in all eukaryotes, is driven by SNARE proteins. Ykt6 is an essential SNARE that plays critical roles throughout the secretory, endocytic, and autophagy pathways. Ykt6 activity is thought to be regulated by a conformational change from a closed cytosolic form to an open membrane-bound form, yet the mechanism that regulates this transition is unknown. Through genetic, pharmacologic, and structural modeling approaches in mammalian cells, we found that phosphorylation regulates Ykt6 conversion from a closed to an open state. The phosphorylation site we identified is highly conserved in evolution and is regulated by the Ca^2+^-dependent phosphatase, calcineurin. We found that phosphorylation is a key determinant for intracellular localization of Ykt6 and its function in macroautophagy. Our studies reveal a novel mechanism by which Ykt6 conformation and activity is regulated by Ca^2+^ signaling with implications in Parkinson’s Disease in which Ykt6 has been shown to play a role.

## INTRODUCTION

Membrane trafficking requires a transport vesicle to reach, dock, and fuse with the correct target membrane. Membrane fusion is commonly considered the final stage in vesicle trafficking and is largely driven by SNARE proteins, which provide the energy for fusion and contribute to the specificity of the trafficking event (Weber et al., 1998) (Han, Pluhackova, & Bockmann, 2017). Ykt6 is an essential R-SNARE and is highly conserved in all eukaryotes. Ykt6 evolutionary conservation is such that human Ykt6 can complement Ykt6-deficient yeast (McNew et al., 1997). Conversely, the yeast Ykt6 can be targeted to the correct intracellular location in mammalian cells (Hasegawa et al., 2003). Ykt6 has been critically involved in vesicular transport between various organelles including the endoplasmic reticulum (ER), Golgi apparatus (Golgi), plasma membrane, and vacuole/lysosome (Gordon et al., 2017) (McNew et al., 1997).

In addition to its roles in the secretory and endocytic pathways, Ykt6 has recently been implicated in macroautophagy (hereafter referred to as autophagy) (Gao, Reggiori, & Ungermann, 2018) (Bas et al., 2018) (Takats et al., 2018) (Matsui et al., 2018). Autophagy is a highly conserved intracellular degradation pathway of long-lived proteins, misfolded proteins, and damaged organelles (Mizushima & Komatsu, 2011). This dynamic process is comprised of three main steps: formation of autophagosomes, fusion of autophagosomes with lysosomes, and lysosomal degradation of autophagic cargo (Zhao & Zhang, 2019). The fusion of autophagosomes with lysosomes is a complex process that involves the cytoskeleton, phosphoinositides, Rab proteins, tethering factors, and SNAREs (Nakamura & Yoshimori, 2017). Key SNAREs involved in this function are Stx7, Stx17, SNAP29, and Vamp7/8. Although recent studies have implicated Ykt6 in autophagosome/lysosome fusion in mammalian cells (Takats et al., 2018) (Matsui et al., 2018), there are discrepancies between specific findings regarding the role Ykt6 plays relative to other SNAREs. To date, mechanisms underlying Ykt6 recruitment to the autophagosome/lysosome interface remain unresolved.

Ykt6 SNARE activity has been proposed to be regulated by lipid modifications that allow Ykt6 to cycle between the cytosol and membrane-bound compartments (Fukasawa, Varlamov, Eng, Sollner, & Rothman, 2004) (Meiringer, Auffarth, Hou, & Ungermann, 2008) (Pylypenko et al., 2008). The C-terminal lipid anchor motif, CCAIM, can be palmitoylated at the first cysteine and farnesylated at the second cysteine (Fukasawa et al., 2004). In addition to the lipid anchor motif, Ykt6 contains an N-terminal regulatory Longin domain followed by the SNARE domain. When Ykt6 is in the cytosol, it has been proposed to be in a closed conformation whereby the N-terminal Longin domain is folded back onto the SNARE domain as revealed by nuclear magnetic resonance (NMR) (Tochio, Tsui, Banfield, & Zhang, 2001). In this case, farnesylation was shown to be an important stabilizer of this closed conformation facilitating hydrophobic interactions between the Longin and SNARE domains (Dai et al., 2016; Wen et al., 2010). It has been suggested that Ykt6 can relocate from the cytosol to membrane-bound compartments by undergoing a large conformational change triggered by the dissociation of the Longin domain from the SNARE domain (Fukasawa et al., 2004) (Wen et al., 2010). This would facilitate its palmitoylation and membrane association. This model is supported by two observations. First, deletion of the Longin domain localizes the protein to both plasma membrane and Golgi (Fukasawa et al., 2004). Second, introduction of a negative charge in F42 (F42E) within the Ykt6 Longin domain which causes the domain to dissociate from the SNARE domain and lead to both Golgi and plasma membrane localization (Fukasawa et al., 2004; Tochio et al., 2001). Because the closed conformation is not membrane-bound and is primarily found in the cytosol, it has been assumed to be inactive. The open conformation, which is membrane-bound, has been postulated to be active. Whether this is a physiologic mechanism that regulates Ykt6 cellular activities *in vivo* remains to be shown.

Here, we report that phosphorylation within the Ykt6 SNARE domain drives an intramolecular change that mediates its conversion from a closed to an open state. We found that the regulation of this conformational change is a key determinant for its intracellular localization and its activities during autophagy. The phosphorylation site we identified is highly conserved throughout evolution and is physiologically regulated by calcineurin, a central Ca^2+^-dependent phosphatase that plays key roles during Ca^2+^ signaling. Taken together, our results provide mechanistic insight into fundamental aspects of Ykt6 regulation and its activities during autophagy. This insight can have implications for Parkinson’s Disease wherein malfunctions in Ykt6 and autophagy have already been implicated (Thayanidhi et al., 2010) (Cooper et al., 2006) (Cerri & Blandini, 2018).

## RESULTS

### A conserved phosphorylation site in the Ykt6 SNARE domain is sensitive to the phosphatase, calcineurin, in both yeast and humans

Our group has established a central role for the 12-kDa cis-trans proline isomerase FK506-binding protein (FKBP12) and the Ca^2+^-dependent phosphatase, calcineurin, in *α*-synuclein (*α*-syn) toxicity, a protein critically implicated in Parkinson’s Disease (Caraveo et al., 2017). Specifically, we have found that *α*-syn leads to a pathological increase in cytosolic Ca^2+^ and an increase in calcineurin/FKBP12 activity which leads to cell death (Caraveo et al., 2014; Caraveo et al., 2017). To identify the substrates dephosphorylated by calcineurin/FKBP12 that could lead to this toxic response, we undertook an unbiased phosphoproteomic approach in a yeast model of D-syn toxicity (Caraveo et al., 2017). We focused on those hypophosphorylated peptides that, in the presence of *α*-syn, gained phosphorylation when knocking out both calcineurin and FKBP12. With these criteria, we retrieved two phosphosites (S176, S178) from Ykt6 (Caraveo et al., 2017) (Figure 1A). Both phosphosites lie within the yeast Ykt6 SNARE domain, but only S176 is highly conserved throughout evolution (Figure 1B,C).

**Figure 1.**
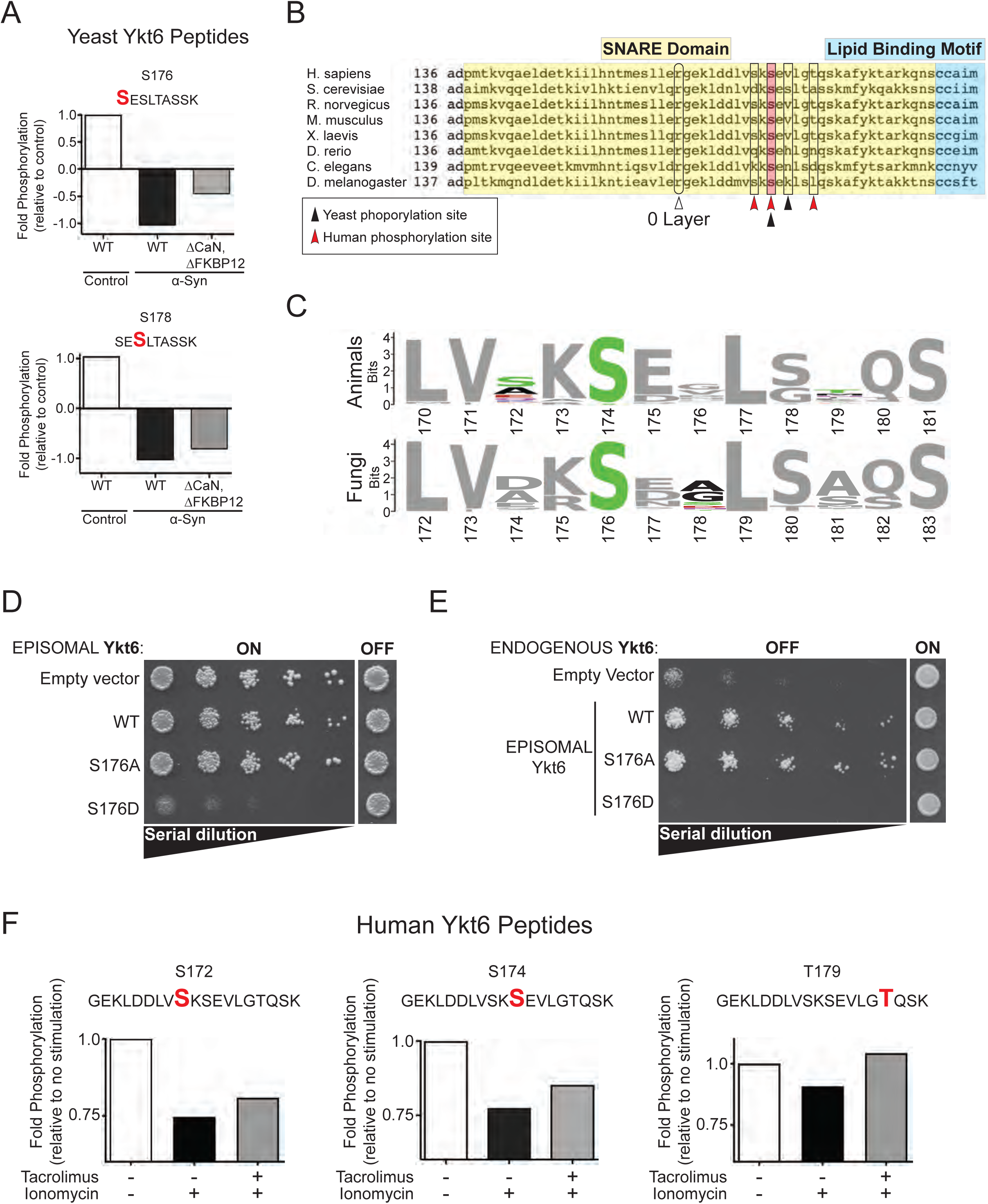
Phosphorylation sites in Ykt6 SNARE domain are sensitive to calcineurin and evolutionarily conserved in yeast and in humans. (A) Fold phosphorylation of the indicated peptides from endogenous yeast Ykt6 detected by shotgun phosphoproteomics after correction for protein abundance from control yeast cells and yeast cells with high levels of Ca^2+^ (driven by overexpression of α-syn) with either WT or knockout for calcineurin (ΔCaN) and knockout for modulator of calcineurin (ΔFKBP12). The identified phosphorylation sites are highlighted in red. Data from triplicate samples was pulled together for illustrated analysis ^28^. **(B)** Alignment of Ykt6 SNARE domain sequences across species. Serines highlighted in pink are the identified phosphorylation site conserved across evolution. Arrows depicted represent Ykt6 0-layer arginine (white) and additional calcineurin-sensitive sites identified in phosphoproteomic screen (red is human; black is yeast). **(C)** Animal and fungal Ykt6 protein sequences by sequence logos obtained from Tracey database and aligned. The residues associated with evolutionary conserved positions are shown. The calcineurin-dependent phosphorylation sites retrieved from the mass spectrometry screens are shown in green. **(D)** Wild type yeast cells were spotted onto plates containing synthetic defined (SD)-Leu media; episomal Ykt6-Leu selective, and replica plated in five-fold serial dilutions onto episomal Ykt6-inducing plates [Galactose (SGal)-Leu; episomal-Leu selective: empty vector, Wild type (WT), phospho-ablative (S176A) and/or phospho-mimetic (S176D)]. Representative plate of n=3. **(E)** Ykt6 temperature sensitive yeast strain were spotted onto plates containing synthetic defined (SD)-Leu media; episomal Ykt6-Leu selective, and replica plated in five-fold serial dilutions onto episomal Ykt6-inducing plates [Galactose (SGal)-Leu; episomal-Leu selective: empty vector, Wild type (WT), phospho-ablative (S176A) and/or phospho-mimetic (S176D)]. Endogenous Ykt6 is depleted by incubating the cells at 37°C, the non-permissive temperature. Representative plate of n=3. **(F)** Fold phosphorylation of the indicated human Ykt6 peptides from HEK293T cells as detected by iTRAQ mass spectrometry. Prior to GFP-Ykt6 immunoprecipitation, cells were treated for 30 minutes with the Ca^2+^ ionophore ionomycin (1μM) and co-treated with calcineurin-specific inhibitor Tacrolimus (1μM).

As a first test of the physiological relevance of the evolutionary conserved phosphorylation site, we overexpressed yeast Ykt6 wild-type (WT), phospho-ablative (S176A), or phospho-mimetic (S176D) mutant under the galactose inducible promoter to test for any changes in viability (Figure 1D). While overexpression of the phospho-ablative mutant had no effect on cell growth, overexpression of the phospho-mimetic mutant compromised the ability of the yeast cells to grow. Similar results were found in a yeast temperature sensitive strain in which the expression of Ykt6 is abolished at the non-permissive temperature 37 °C (Kofoed et al., 2015). In these yeast, the lack of Ykt6 results in a growth defect that can be rescued by episomal expression of WT Ykt6 (Figure 1E). If the conserved phosphorylation site plays an important role in Ykt6 function, we reasoned that mutations on this site might fail to rescue viability. Expression of the phospho-ablative mutant was able to complement cell growth similar to WT; however, expression of the phospho-mimetic mutant did not rescue (Figure 1E). These results indicate an important biological role of the evolutionary conserved calcineurin–dependent phosphosite in yeast.

We next investigated whether human Ykt6 is similarly phosphorylated and sensitive to calcineurin phosphatase activity. We transfected human embryonic kidney (HEK293T) cells with human Ykt6 N-terminally tagged with green fluorescent protein (GFP). Importantly, N-terminal tagging of Ykt6 does not affect its function as shown by other groups (Fukasawa et al., 2004). To mimic the high calcineurin activity from *α*-syn expression in yeast, transfected cells were treated with the Ca^2+^ ionophore, ionomycin, to trigger a rapid increase in cytosolic Ca^2+^ and hence calcineurin activation. In other preparations, cells were co-treated with ionomycin and the calcineurin-specific inhibitor, Tacrolimus. Ykt6 was immunoprecipitated from each condition and subjected to mass spectrometry by isobaric tag for relative and absolute quantitation (iTRAQ) across treatments. We retrieved three phosphosites in human Ykt6 (S172, S174, T179) whose phosphorylation was decreased in the presence of ionomycin and was partially restored with the addition of Tacrolimus (Figure 1F). Importantly, one of the phosphosites identified in human Ykt6 (S174) is homologous to the conserved site (S176) retrieved from the yeast phosphoproteomic screen (Figure 1B-C). Together, these results indicate that the phosphorylation site we identified in the Ykt6 SNARE domain is both evolutionarily conserved and calcineurin-sensitive in yeast and in humans, prompting further investigation.

### Calcineurin-sensitive phosphorylation site in human Ykt6 SNARE domain is a key determinant for its intracellular localization

To examine if the evolutionarily conserved calcineurin-sensitive phosphosite is physiologically relevant in mammalian cells, we overexpressed N-terminal GFP fusions of human wild-type (WT), phospho-ablative (S174A), or phospho-mimetic (S174D) mutant Ykt6 in HEK293T and HeLa cells. Compared with each other, the mutants showed marked differences in their intracellular localization (Figure 2A). We found no difference in their transient expression levels (Supplemental Figure 2A-B). Of note, the phospho-ablative mutant resembled the WT in its cytosolic localization, whereas the phospho-mimetic mutant mainly localized to the plasma membrane. Consistent with this result, cell fractionation demonstrated that the phospho-mimetic mutant was increased in the membrane fraction relative to both WT and the phospho-ablative mutant (Figure 2B-C and Supplemental Figure 2C). We next examined their co-localization with ER and trans-Golgi markers. Both phosphomutants showed strong co-localization with the trans-Golgi marker (Figure 2D-E) and to a lesser extent with the ER marker (Figure 2F-G).

**Figure 2.**
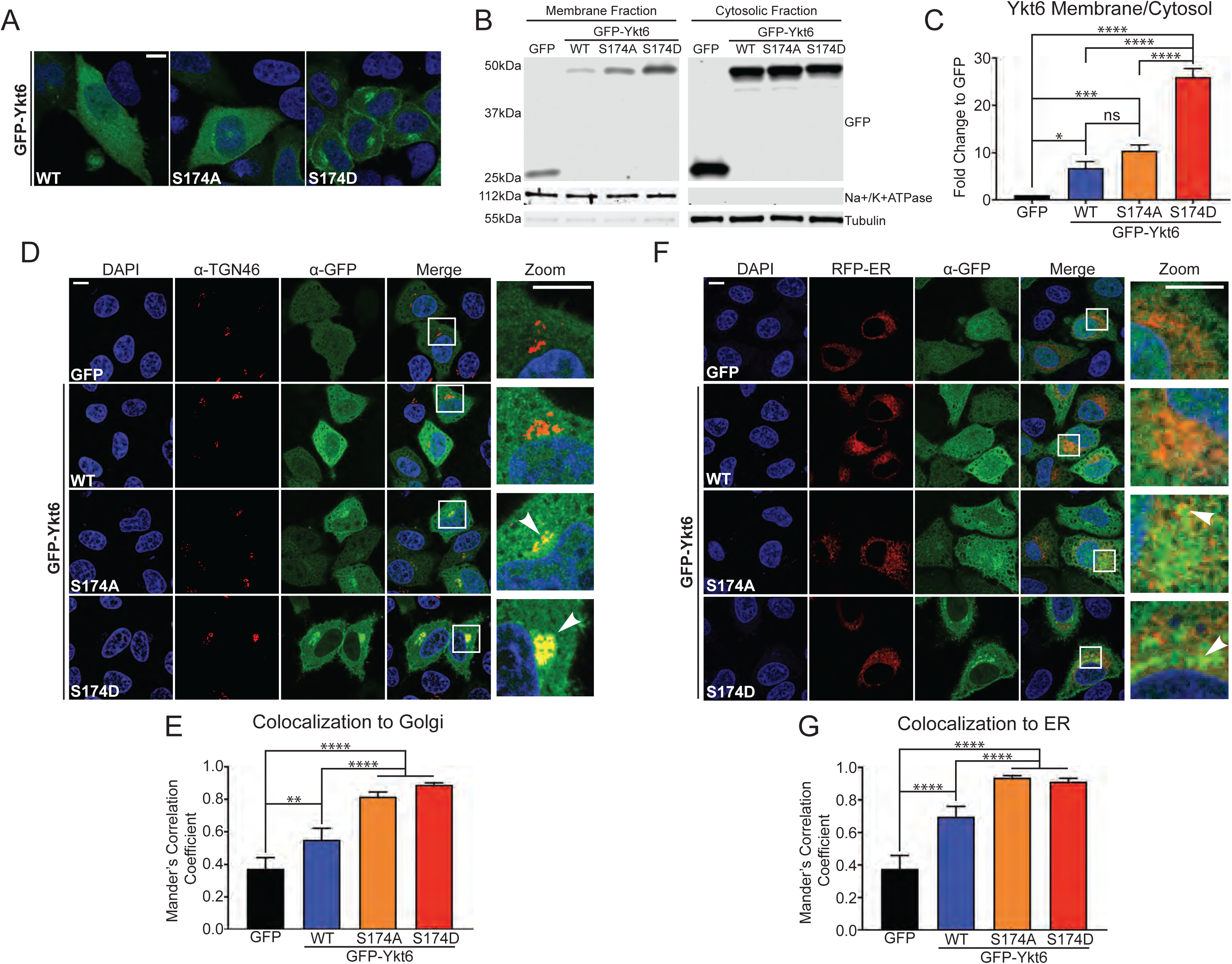
Evolutionarily conserved phosphorylation site within human Ykt6 SNARE domain (S174) is a critical determinant for its intracellular localization. (A) Representative immunofluorescence images of transiently transfected HeLa cells with GFP, GFP-tagged wild-type (WT), phospho-ablative mutant (S174A) and phospho-mimetic mutant (S174D) of Ykt6. Nuclei (blue) are stained with DAPI. **(B)** Representative Western Blot for membrane and cytosolic fractions of HEK293T cells transiently transfected as described in (A). Membrane (Na+/K+ ATPase) and cytosolic (tubulin) markers serve as controls for fractionation purity. **(C)** Fold change of membrane/cytosol fraction calculated as: 1) normalizing to GFP/actin (see supplemental Figure 2C), 2) obtaining the ratios of GFP-Ykt6 fusion protein to tubulin (for the cytosolic fraction), GFP-Ykt6 fusion protein to Na+/K+ ATPase (for the membrane fraction) and 3) membrane fraction/cytosolic fraction. N=4 *p,0.05, **p<0.01, ***p,0.001 One-way ANOVA, Fisher’s uncorrected LSD test. **(D)** Representative immunofluorescence images of transiently transfected HeLa cells with GFP, GFP-tagged WT or phosphomutants of Ykt6 and immunostained for TGN-46, a trans-Golgi marker. Nuclei (blue) are stained with DAPI. White arrows point to regions of colocalization. **(E)** Colocalization analysis based on Mander’s correlation coefficient for TGN-46 and Ykt6. **(F)** Representative immunofluorescence images of transiently co-transfected HeLa cells expressing GFP, GFP-tagged wild-type (WT) or indicated phosphomutants of Ykt6 along with ER-RFP to delineate the ER. White arrows point to regions of colocalization. Nuclei (blue) are stained with DAPI. **(G)** Colocalization analysis based on Mander’s correlation coefficient between ER and Ykt6. Scale bars in (A,D,F) are 10μm. N=3 **p<0.01 ****p<0.0001 One-way ANOVA, Tukey’s test.

To investigate whether the plasma membrane localization of the phospho-mimetic mutant is dependent on calcineurin, we took a pharmacologic approach. HeLa cells which were transfected with N-terminal GFP fusions of WT, the phospho-ablative, or the phospho-mimetic mutant. These cells were Ca^2+^ stimulated with ionomycin and/or treated with the calcineurin-specific inhibitor, Tacrolimus. If the phosphorylation site (S174) is sensitive to calcineurin, we reasoned that inhibition of calcineurin should relocalize WT Ykt6 from the cytosol to the plasma membrane. Conversely, the intracellular localization of both the phospho-mimetic and phospho-ablative mutants should not be affected by calcineurin inhibition. Indeed, we found that inhibition of calcineurin caused an increased in WT Ykt6 plasma membrane localization (Figure 3A-B). Inhibition of calcineurin under Ca^2+^ stimulation by ionomycin further increased WT Ykt6 plasma membrane localization, suggesting that phosphorylation at this site is also Ca^2+^-dependent.

**Figure 3.**
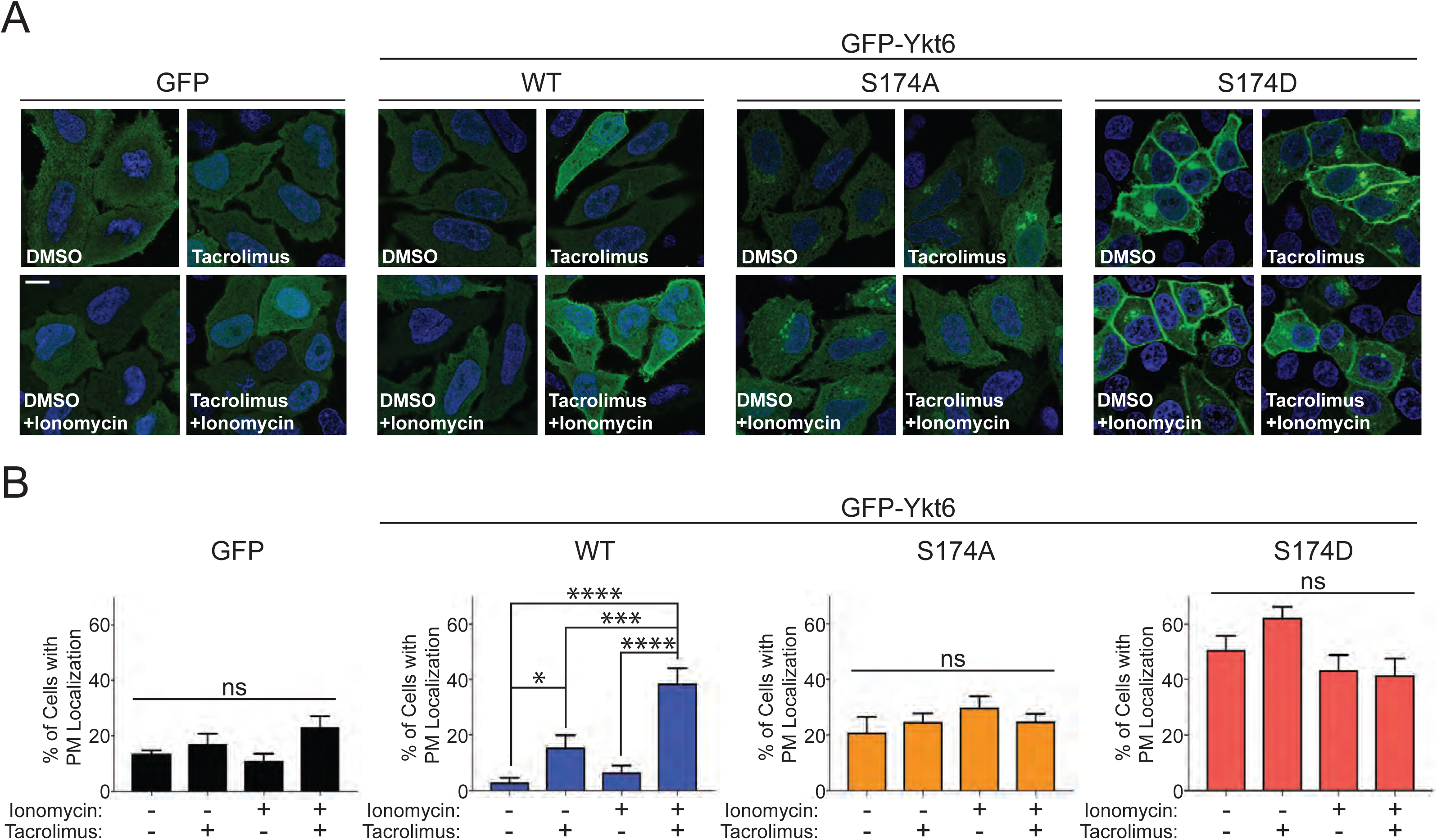
The intracellular localization of Ykt6 is dependent on calcineurin activity. (A) Representative immunofluorescence images of transiently transfected HeLa cells with GFP-tagged wild-type (WT) or indicated phosphomutants of Ykt6. Cells were treated with ionomycin (1μM) and/or Tacrolimus (1μM) for 30 minutes. Nuclei (blue) are stained with DAPI. Scale bar is 10μm. **(B)** Quantification of cells with GFP plasma membrane localization as shown in (A). N=3 *p<0.05 ***p<0.001 ****p<0.0001 One-way ANOVA, uncorrected Fisher’s LSD test.

Importantly, we did not observe any significant changes in intracellular localization in cells transfected with GFP alone, the phospho-ablative, or the phospho-mimetic mutant under the same pharmacological treatment. Together, these results indicate that the conserved phosphosite (S174) within the human Ykt6 SNARE domain is regulated by calcineurin and is a critical determinant for Ykt6 plasma membrane localization.

### Phosphorylation of the evolutionarily conserved site within the SNARE domain regulates Ykt6 conformation

Previous work on Ykt6 demonstrated that a point mutation within the Ykt6 Longin domain (F42E) could drive the protein to its open conformation (Fukasawa et al., 2004). In addition, the F42E mutant had an altered intracellular localization pattern versus control (i.e., from the cytosol to the plasma membrane and Golgi). A similar intracellular localization was found in a deletion mutant of the entire regulatory Longin domain (ΔN) (Tochio et al., 2001). These mutants suggest a model in which the extent of interactions between the Longin and SNARE domains is a key determinant in the conformational state and intracellular localization of Ykt6. Interestingly, the localization pattern of the phospho-mimetic Ykt6 mutant (Figures 2-3) is strikingly similar to that reported for the ΔN or F42E mutant. We wondered whether phosphorylation of S174 in the SNARE domain induces a conformational change in Ykt6.

To directly examine the effect of S174 phosphorylation on Ykt6 conformation, we used partial proteolysis with trypsin. This method can assess protein conformation by subjecting purified proteins to different concentrations of the protease trypsin for a given amount of time. For proteins in a closed, compact conformation, higher concentrations of trypsin are required to access all the available arginine and/or lysine residues. Therefore, these proteins will produce less fragments when run on an SDS gel after trypsin digest. On the other hand, for proteins in a more open, less compact conformation, lower concentrations of trypsin are sufficient to access all available arginine and/or lysine residues. Hence, these proteins will produce more fragments. We purified GFP-tagged WT Ykt6 and its phosphomutants from HEK293T cells using GFP antibodies (Figure 4A and Supplemental Figure 4A,B) and subsequently subjected the purified proteins with different concentrations of trypsin for 1 hour. The WT and phospho-ablative mutant displayed their first cleavage product (∼15 kDa) with 10 ng of trypsin. This was accompanied by a concomitant disappearance of the larger molecular weight band (∼ 50 kDa, see top arrow in Figure 4A). The phospho-mimetic mutant, however, displayed its first and second cleavage products with 5 ng of trypsin (Figure 4A,B). This cleaved band was specific to Ykt6, since it did not react when probed with GFP, the N-terminal fusion tag we used for purification (Supplemental Figure 4A). These data suggest that the phospho-mimetic mutant is less compact and more open as compared to the phospho-ablative mutant and WT Ykt6.

**Figure 4.**
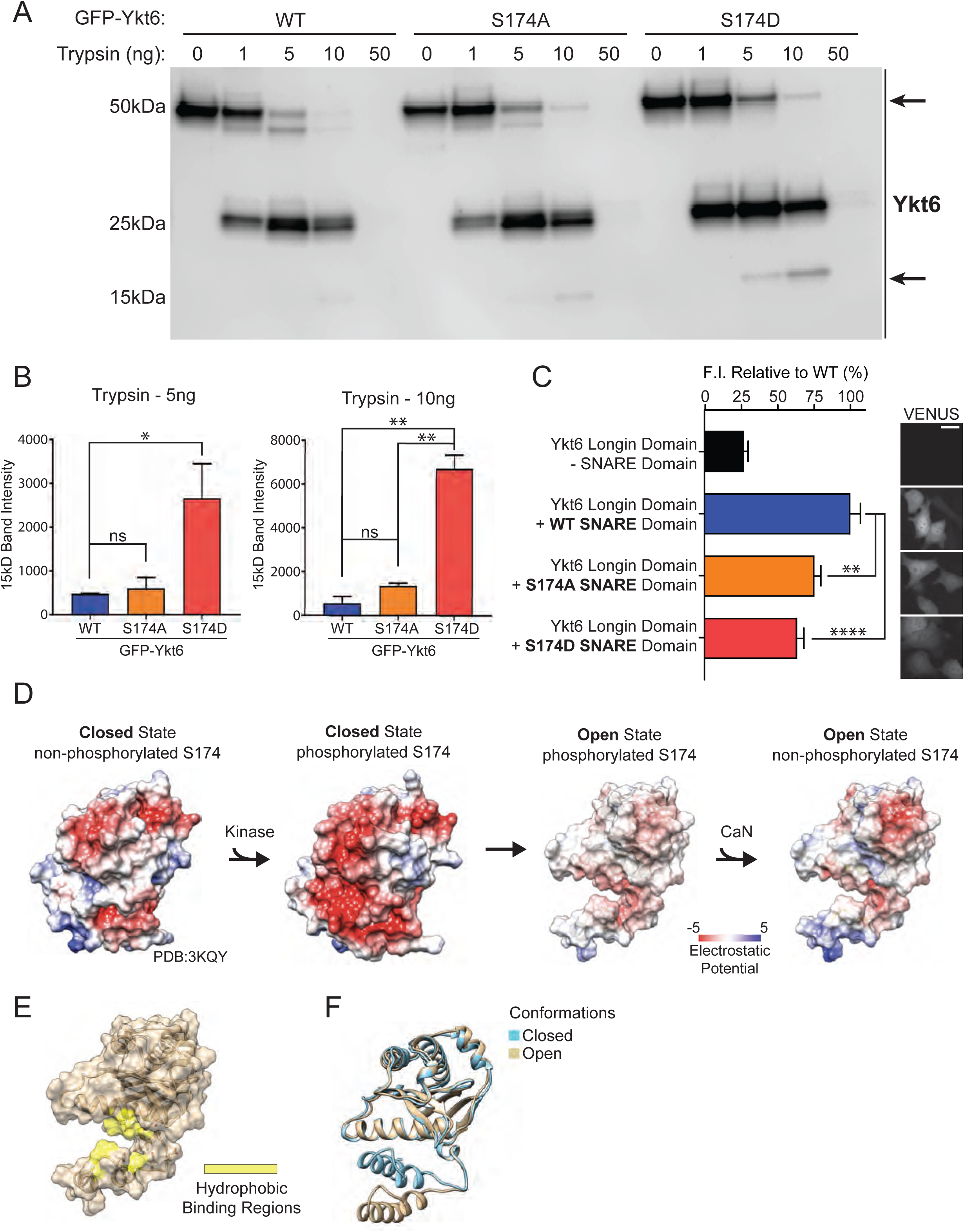
Phosphorylation within the evolutionarily conserved site in Ykt6 SNARE domain regulates its conformation. **(A)** Representative western blot for Ykt6 from GFP-purified WT and phosphomutants of human Ykt6 incubated with the indicated amounts of trypsin for 1 hour at 25°C. Arrows indicate specific cleavage products. **(B)** Densitometry analysis of the 15 kDa cleavage band. N=3 *p<0.05 *p<0.01 One-Way ANOVA, Tukey’s test. **(C)** Representative live-cell fluorescence images of HeLa cells transiently expressing indicated combinations of split Venus constructs. Quantification of fluorescence intensity of Venus normalized by the area of the cell. Wild-type (WT) Longin domain with the SNARE domains of either: wild-type (SNARE WT), phosphoablative (SNARE S174A) or phosphomimetic (SNARE S174D) of human Ykt6. Scale bar is 10μm. N=4 * p<0.05 One-Way ANOVA, Tukey’s test. **(D)** Electrostatic potential computed and projected at the surface representation of both, open and closed conformation, in the absence and presence of phosphorylation at the evolutionarily conserved site S174 (pS174). Ykt6 exists in a closed conformation and phosphorylation at S174 by a kinase increases the elecrtrostatic potential driving its open conformation. Subsequently, dephosphorylation by calcineurin (CaN), decreases its electrostatic potential. **(E)** Surface model of Ykt6 open conformation depicting the residues mediating hydrophobic binding in yellow when in the closed conformation. The cluster of negatively charged residues is shown as sticks. **(F)** Overlap of Ykt6 open and closed conformation.

To further test the hypothesis that phosphorylation within Ykt6 SNARE domain can prevent the interaction between the SNARE and the Longin domains, we used a fluorescence complementation approach in HeLa cells. One vector contained the N-terminal half of Venus fused to Ykt6 Longin domain, and the second vector contained the C-terminal half of Venus fused to Ykt6 SNARE domain of either WT, phospho-ablative, or phospho-mimetic mutants. All these constructs lacked the lipid anchor motif CCAIM to avoid complications due the inherent differences in their subcellular localization. If phosphorylation is a driver of Ykt6 open conformation, we would expect the phospho-mimetic mutant to have less complementation between the split Venus domains compared to the phospho-ablative mutant and/or the WT. This change would be manifested as decrease in Venus fluorescence. Expression of a single split Venus fusion construct had little to no background fluorescence (Figure 4C and Supplemental Figure 4C), however co-expression of the WT SNARE domain with the Longin domain generated a robust Venus fluorescent signal indicative of an interaction between the Longin and SNARE domains (Figure 4C). While Venus fluorescence was reduced by the phospho-ablative SNARE domain, the phospho-mimetic SNARE domain showed the least Venus fluorescence. This experiment shows that the presence of a phosphate group at S174 can reduce the interaction between the SNARE domain and the Longin domain and suggests a possible role of this phosphorylation site in promoting Ykt6 open conformation.

To understand how phosphorylation at S174 impacts Ykt6 open conformation we took a bioinformatic structural approach (Figure 4D-F). It is known that Ykt6 closed conformation is stabilized by an extended network of hydrophobic amino acids in the Longin domain (such as F34, F42) and in the SNARE domain (such as V171), which when individually mutated to a glutamate leads to an open conformation (Wen et al., 2010). The S174 site is located at the end of the *α*F helix in the SNARE domain which makes it accessible to phosphatases and kinases (Supplemental Figure 4D-E). When we modeled the phosphorylated form of this residue using the available Ykt6 closed conformation (Wen et al., 2010), the S174 phosphate group could be placed into the structure without introducing unresolvable clashes of atoms due to spatial constraints (Supplemental Figure 4E). Given that electrostatic interactions can significantly influence protein conformation, we searched for negatively charged regions that, in addition to our identified phosphorylated sites, could contribute to the repulsion of the Longin and SNARE domains. We noted a conserved patch of negatively charged residues in close proximity to our calcineurin-dependent phosphorylation sites in the *α*F helix SNARE domain, and at the Longin domain around a short helix turn (*α*E) (Figure 4E and Supplemental Figure 4F). To investigate whether the highly conserved negatively charged patch might contribute to an electrostatic repulsion between the helices, we slightly adjusted the Ykt6 closed conformation. We did so by fine tuning the torsion angles of the loop region, residues 162 to 164. This allows visualization of the compact electrostatic potential in this region that would otherwise be visible only at the surface. After such adjustment, it became clear that the electrostatic repulsion between these regions could drive displacement of the Longin and SNARE domain and lead to Ykt6 open conformation (Figure 4D-F). Thus, our data suggests that electrostatic repulsion acting upon longer distances prevents short-range hydrophobic binding and therefore favors the open conformation. Taken together, the trypsin digestion, the biomolecular fluorescence complementation experiments, and the structural modeling, supports a critical role of phosphorylation at the evolutionarily conserved site in Ykt6 SNARE domain in destabilizing the hydrophobic interaction between the Longin and SNARE domains, promoting the open conformation.

### Phosphorylation at the evolutionarily conserved calcineurin-dependent site regulates human Ykt6 function in autophagy

Ykt6 plays an important role in multiple steps of the secretory, endocytic and autophagy-lysosomal pathways. The mechanisms that regulate these distinct activities are not well-understood. We have found that relative to the WT, the human Ykt6 phospho-mimetic mutant has a distinctive plasma membrane, trans-Golgi and to a lesser extent ER localization (Figure 2D-G). The phospho-ablative S174A mutant, however, is not associated with the plasma membrane, but it is also localized with the trans-Golgi and ER (Figure 2D-G). If Ca^2+^-calcineurin activation is a key signaling switch that enables Ykt6 conformational change and intracellular localization, this should affect Ykt6 cellular activities and therefore, the spectrum of its interactors.

To test this hypothesis, we took a mass spectrometry approach. GFP immunoprecipitants from HEK293T cells transfected with N-terminal GFP-tagged fusions of WT human Ykt6 and the corresponding phospho-ablative and phospho-mimetic mutants were subjected to three independent rounds of mass spectrometry. Our immunoprecipitations were robust, specific to Ykt6 and importantly, very similar across WT and the Ykt6 phosphomutants (Supplemental Figure 5). Hits were scored as positive based on the following criteria: 1) present in three mass spectrometry analyses, 2) spectral counts had to be at least 5 for either Ykt6 WT and/or phosphomutants if the GFP control pulldown contained zero spectral counts, and/or 3) spectral counts had to be at least 10-fold higher for Ykt6 WT and/or phosphomutants if the GFP pulldown control contained greater than zero spectral counts. Under these criteria we retrieved a total of 15 Ykt6 specific interactions (Figure 5A,B and Supplemental Table 1). Eleven protein interactions were common between WT Ykt6 and phosphomutants (Figure 5A,B), however the ratios of these interactions changed depending on the phosphorylated state (Figure 5A and Supplemental Table 1). To understand how the common hits differ between WT and phosphomutants, we normalized the spectral counts to WT Ykt6. In most cases, the phospho-ablative mutant had increased protein interactions with SNAREs compared to both the WT and the phospho-mimetic mutant. These included some of the already established interacting partners of Ykt6 in vesicular trafficking such as GOSR2, Sly1 (SCFD1 gene) and in autophagy such as SNAP29 and Stx17 (Figure 5A and Supplemental Table 1). In most cases, the phospho-mimetic mutant resembled the WT in the magnitude of interactions with the SNAREs SNAP47, Stx8, VTI1B and other already known interacting partners such as GOSR2 and Sly1 (Figure 5A and Supplemental Table 1). The phospho-mimetic mutant showed increased interactions relative to the WT but lower than the phospho-ablative mutant which included SNAP29, Stx18, Snap-*α* (NAPA gene) and NBAS In some cases, the ratios of the interacting partners between the phospho-mimetic and the phospho-ablative mutant did not change; these included the interactions with the vesicle fusion ATPase NSF (Figure 5A and Supplemental Table 1). The interactions with the E3 ubiquitin ligase FEM1B and the farnesyl transferases FNTA and FNTB, were the only interactions that did not change between the WT and phosphomutants (Figure 5A and Supplemental Table 1). Furthermore, we detected a few unique interactions gained in the phosphomutants. The phospho-ablative mutant interacted specifically with Stx17, a protein critically involved in facilitating autophagosome/lysosome fusion during autophagy (Itakura, Kishi-Itakura, & Mizushima, 2012) (Figure 5A and Supplemental Table 1). The phospho-mimetic Ykt6 mutant on the other hand, lost the interaction with Beta Galactosidase GLB1, a lysosomal enzyme genetically linked to GM1-gangliosidosis (Figure 5A and Supplemental Table 1).

**Figure 5.**
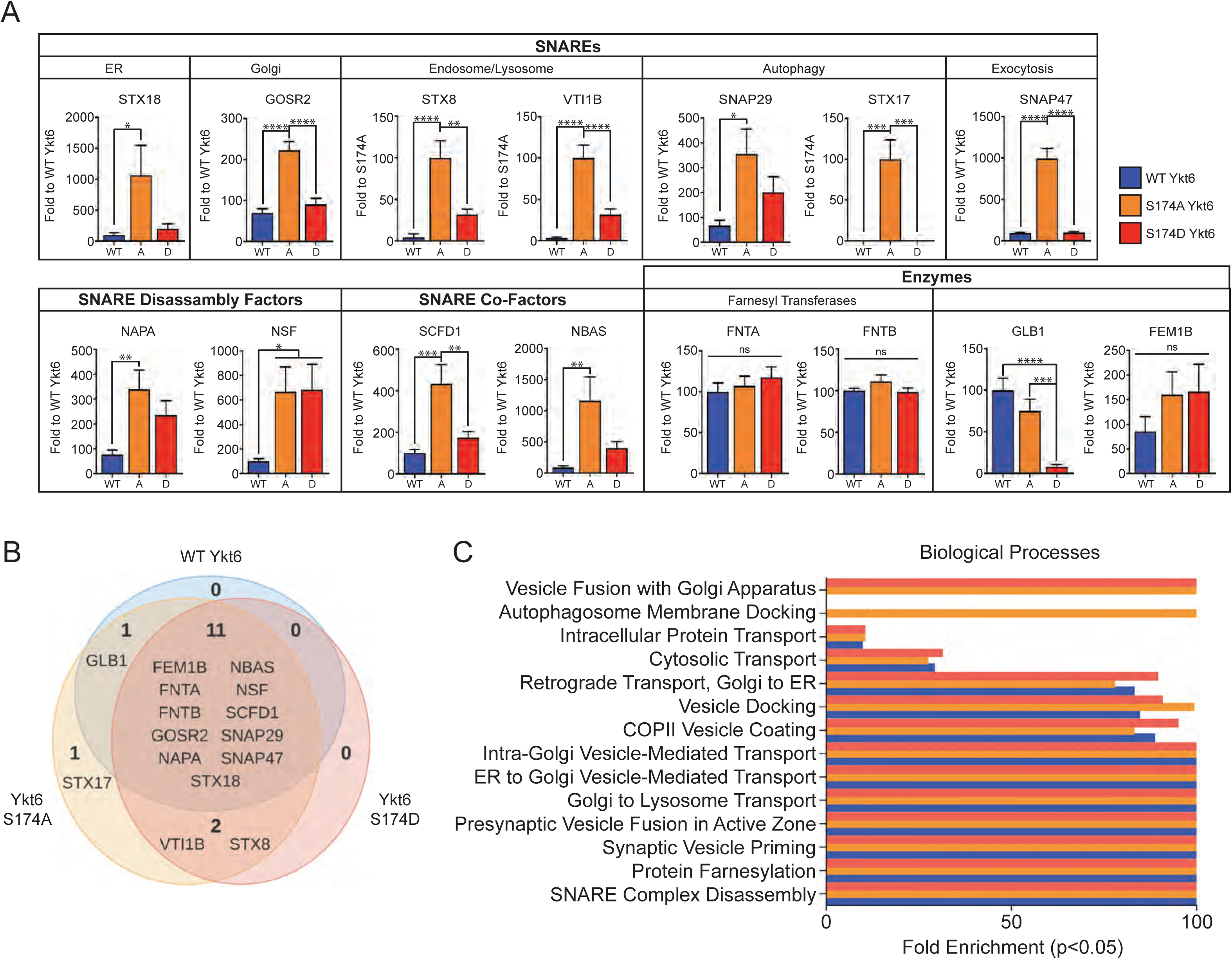
Phosphorylation of the evolutionarily conserved site in Ykt6 SNARE domain S174 affects the specificity of its binding partners. (A) Quantitative analysis of mass spectrometry hits that passed the selection criteria described in main text and methods. Spectral counts for common interactors were normalized to WT Ykt6. N=3 independent mass spectrometry runs. *p<0.05 **p<0.01 ***p<0.001 ****p<0.0001 One-way ANOVA, Tukey’s test. **(B)** Venn diagram representation of all mass spectrometry hits that passed the selection criteria described in main text and methods. **(C)** Gene ontology analysis for biological processes from the mass spectrometry hits in (A).

To understand how the proteins retrieved form the mass spectrometry analysis reflects Ykt6 cellular activities and how these could be perturbed by the phosphomutants, we analyzed the hits at a functional level. We saw an enrichment in Ykt6 function in different vesicular trafficking steps such as the Golgi and ER transport as other groups have reported, but importantly, it pointed to specific cellular pathways whereby Ykt6 phosphorylation might play a key role. The biological processes that were more affected by the phospho-ablative mutant were autophagosome-membrane docking, intra-Golgi transport and Golgi-organization (Figure 5C). In sum, this mass spectrometry analysis revealed three things: 1) Ykt6 can participate in diverse cellular pathways, 2) the affinity of Ykt6 to its binding partners depends on the calcineurin-dependent phosphorylation site S174, and 3) the phosphomutants might play a distinctive role in specific cellular pathways.

Ykt6 has been recently shown to be implicated in two cognate SNARE complexes in autophagosomal/lysosomal fusion (Matsui et al., 2018; Takats et al., 2018). One complex consists of Stx7 (Qa SNARE localized in the lysosome) and SNAP29 (Qbc SNARE) (Matsui et al., 2018). Another complex in which Ykt6 has been implicated consists of Stx17 (Qa SNARE localized in the autophagosome) and SNAP29 (Takats et al., 2018). Our mass spectrometry results showed that the Ykt6 phospho-ablative mutant interacts specifically with Stx17 and SNAP29, while the phospho-mimetic mutant lacks the ability to interact with Stx17 and has decreased binding affinity with SNAP29 (Figure 5A,B and Supplemental Table 1). These findings indicate that the calcineurin-dependent phosphorylation site S174 in Ykt6 may play an active role in regulating the late phases of autophagy such as autophagosome/lysosome fusion.

To confirm our mass spectrometry results, we first took a biochemical approach (Figure 6A,B). N-terminal GFP-tagged fusions of WT, phospho-ablative, or phospho-mimetic human Ykt6 were expressed in HEK293T cells, immunoprecipitated with GFP and immunoblotted for SNAP29 and Stx17. In agreement with the mass spectrometry data, we detected increased binding between the Ykt6 phospho-ablative mutant and SNAP29 and Stx17 compared to the phospho-mimetic mutant and the WT (Figure 6A,B). We observed the same effect when we challenged the cells by starvation. The interactions between Ykt6 phospho-ablative mutant and SNAP29 and Stx17 remained as high as under growing conditions (Figure 6A,B). In contrast, the phospho-mimetic mutant and the WT exhibited very little interaction with SNAP29 and Stx17 under both growing and starvation conditions (Figure 6A,B).

**Figure 6.**
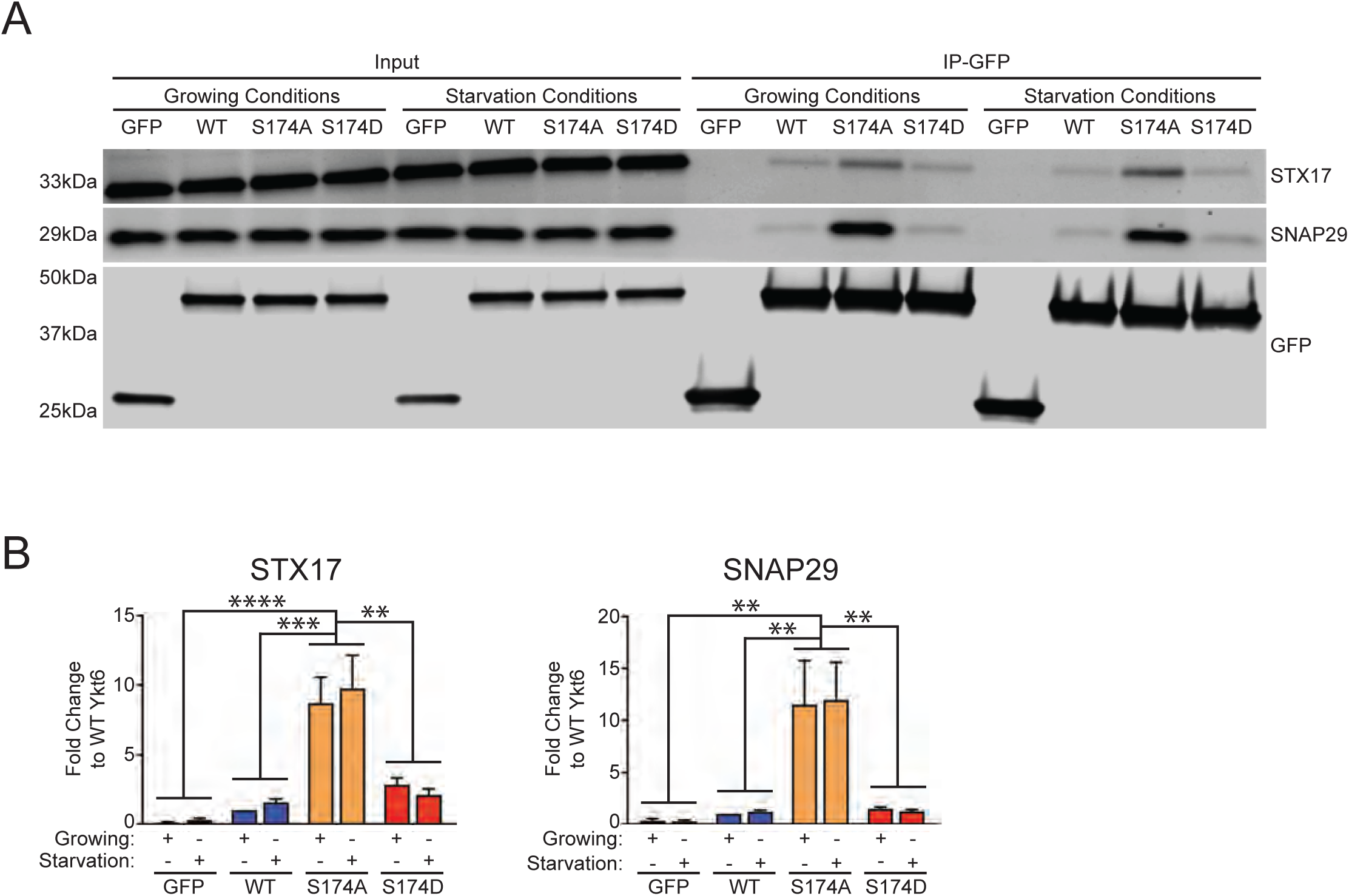
Ykt6 binding affinity for STX17, SNAP29 and Vamp8 is dependent on the phosphorylation at the evolutionarily conserved site in the SNARE domain. (A) Representative western blot of GFP immunoprecipitations from HEK293T cells expressing either GFP, wild-type (WT) GFP-Ykt6 or GFP-Ykt6 phosphomutants. Cells were treated for 2 hours (2h) in growing conditions (fresh 10% FBS and 4.5% glucose growth medium) or in starvation conditions (1X HBSS supplemented with 250nM Torin-1 and 10mM HEPES). **(B)** Quantification of STX17 and SNAP29 (A) relative to the efficiency of the pull down from each condition and normalized to WT Ykt6. N=3 *p<0.05 **p<0.01 ***p<0.001 ****p<0.0001 Two-way ANOVA, uncorrected Fisher’s LSD test.

To investigate whether the differences in binding between the phospho-ablative and the phospho-mimetic mutant with the members of the autophagosome/lysosome fusion machinery could play a functional role in autophagy, we examined the effect of these mutants under starvation. N-terminal GFP-tagged fusions of WT, phospho-ablative, or phospho-mimetic human Ykt6 were expressed in HEK293T cells and grown in complete media (growing conditions) or under starvation conditions with Torin-1, an mTOR inhibitor for 2 hours (Figure 7A-C). To monitor autophagic flux, we examined the expression of the autophagy marker LC3 in the presence and absence of Bafilomycin A_1_ (Baf-A_1_), a potent inhibitor of the vacuolar H^+^ ATPase. The presence of Baf-A_1_ indicates whether there is an increase or decrease in the autophagic flux. No change in LC3-II expression or puncta indicates an inhibition in autophagy whereas an increase in LC3-II expression or puncta indicates an increase in autophagy. By western blot, induction of autophagy increases the expression of lipidated LC3 (LC3II), which migrates at a lower molecular weight band on an SDS gel. By immunofluorescence, induction of autophagy results in an increase in LC3-II puncta. Under normal growing conditions, we did not observe an effect of WT and phosphomutants on basal LC3-II or LC3-II autophagic flux either by western blot or immunofluorescence (Figure 7A,B,E and Supplemental Figure 7A,C). However, under starvation conditions, we observed a block in autophagic flux only in the phospho-mimetic mutant manifested by a lack of increase in LC3-II expression and puncta upon Baf-A_1_ treatment (Figure 7A,C-F and Supplemental Figure 7B-E). These results indicate that the phospho-ablative mutant facilitates autophagosome/lysosome fusion by binding to SNAP29 and Stx17 complex, whereas the phospho-mimetic mutant inhibits autophagosome/lysosome fusion due to lack of interaction with these autophagy SNAREs.

**Figure 7.**
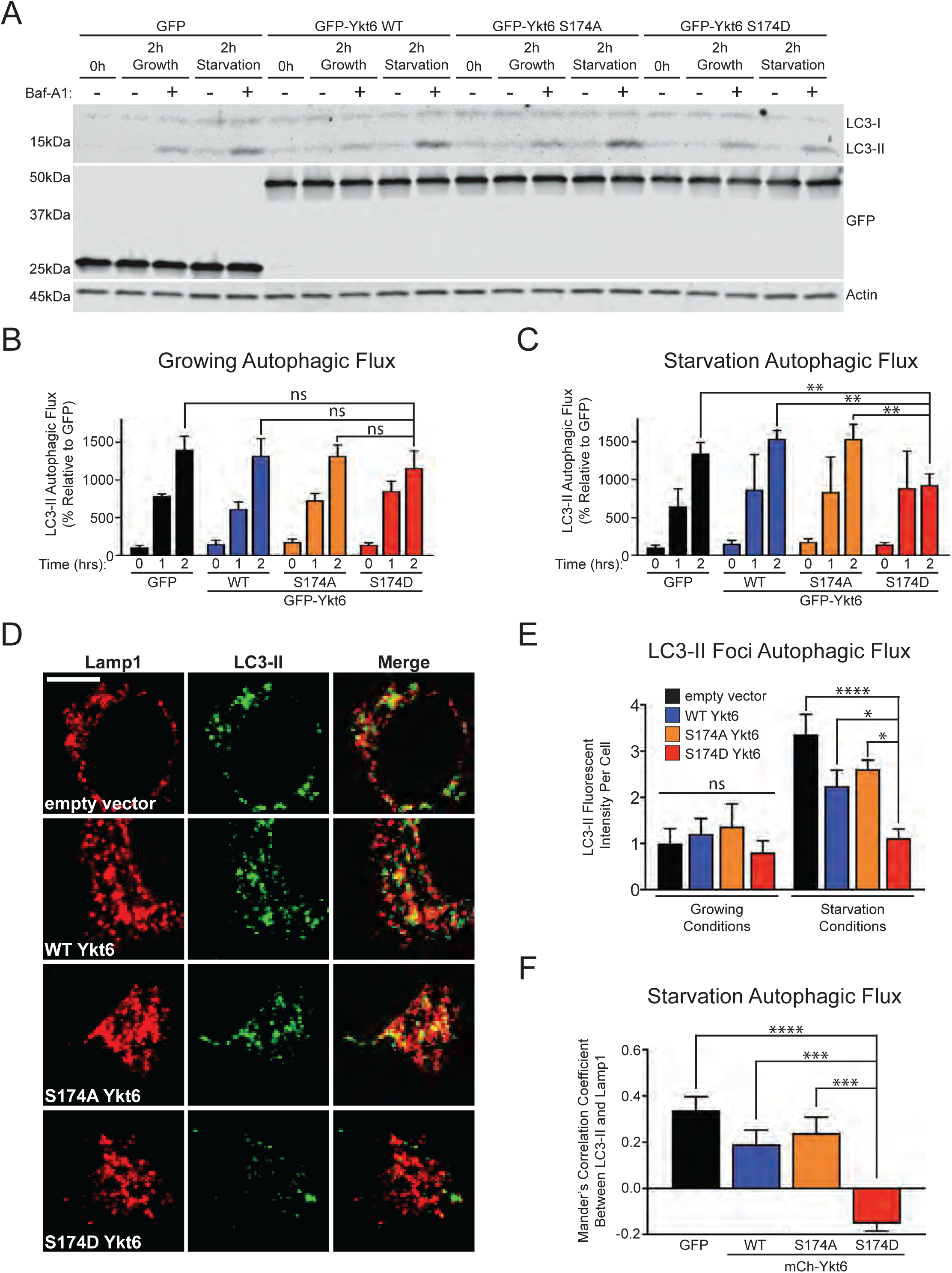
Ykt6 effects on autophagosome/lysosome fusion depend on the phosphorylation at the evolutionarily conserved site in the SNARE domain. Representative western blot for LC3-II from HEK293T transiently expressing GFP-tagged wild-type (WT) Ykt6 and phosphomutants. Cells were treated for 2 hours (2h) with 200nM Bafilomycin A_1_ (Baf-A_1_) in growing conditions (DMEM with 10% FBS and 4.5% glucose growth medium) or in starvation conditions (1X HBSS with 250nM Torin-1 and 10mM HEPES). Actin serves as a loading control. **(B-C)** Autophagic flux (defined as [LC3-II Baf-A_1_] – [LC3-II]) from western blot in (A) of **(B)** growing and **(C)** starvation conditions. (A-C) N=6 *p<0.05 **p<0.01 ****p<0.0001 Two-way ANOVA, Fisher’s uncorrected LSD test. **(D)** Representative immunofluorescence images of overexpressed GFP-LC3 and endogenous Lamp1 colocalization from HeLa cells expressing WT human Ykt6 and phosphomutants, under starvation conditions in the presence of Baf-A_1_ as described above. Scale bar is 5μm. N=3 **(E)** Quantification of (D) autophagic flux from the fluorescence intensity of GFP-LC3-II foci. **(F)** Colocalization analysis of (D) based on Mander’s correlation coefficient between Lamp1 and LC3-II of autophagic flux. (E-F) *p<0.05 **p<0.01 ***p<0.001 ****p<0.0001 One-way ANOVA, Tukey’s test.

There is a possibility that the autophagy effects of the Ykt6 calcineurin-sensitive phosphorylation site are a consequence of deregulations in the secretory pathway in which Ykt6 has also been implicated. To test this we turned to a mammalian ligand-inducible reporter cell line (Gordon, Bond, Sahlender, & Peden, 2010). This system utilizes a ligand-reversible dimer as a reporter of secretion consisting of a mutated FKBP protein fused to GFP. The transport of the fusion protein can be quantitatively monitored using flow cytometry. When these proteins dimerize, they can form large aggregates that are retained in the ER which results in an increase in fluorescence intensity by flow cytometry. Upon the addition of an analog of rapamycin, these aggregates can be efficiently and rapidly secreted from the cells, which results in a drop-in fluorescence intensity over time. We utilized the same conditions as for autophagy (transient transfection and the assay was carried out 24hrs post-transfection) except that in this case we utilized mCherry as control and N-terminal mCherry fusion constructs of Ykt6. We saw no significant difference between the WT and Ykt6 phosphomutants in this cell line (Supplemental Figure 7F). This data suggests that under these conditions, the effects of the phosphorylation site we observe are specific to autophagy and are not due to a consequence in malfunctions in the secretory pathway. These data however, does not rule out a possible role of Ykt6 phosphorylation in the secretory pathway as indicated by our mass spectrometry data. The latter could be more sensitive to Ykt6 concentration, time of Ykt6 exposure and/or cell type-dependent.

Even though HEK293T and HeLa cells have very low levels of endogenous Ykt6 compared to other cell lines such as PC12 and/or primary neurons (Supplemental Figure 7G), they still have endogenous Ykt6. To avoid possible confounding issues due to overexpression, we took a knockdown approach. To do so, we switched to the rat pheochromocytoma cell line, PC12, for two important reasons: 1) endogenous Ykt6 is highly abundant and therefore better suited to test the effects of knockdown, and 2) given that this cell line is rat we can exploit the species differences and knockdown the endogenous rat Ykt6 while rescuing with Ykt6 human constructs. In congruence with our overexpression results from Figure 7, under normal growing conditions, we did not observe an effect of WT Ykt6 or phosphomutants on LC3-II autophagic flux by western blot (Figure 8A-B). However, also in congruence with our overexpression experiments, we observed a block in autophagic flux only with the phospho-mimetic mutant under starvation conditions (Figure 8A,C). Together, these data suggest that phosphorylation at the evolutionary conserved calcineurin-sensitive site in Ykt6 SNARE domain plays an important regulatory role during autophagosome/lysosome fusion.

**Figure 8.**
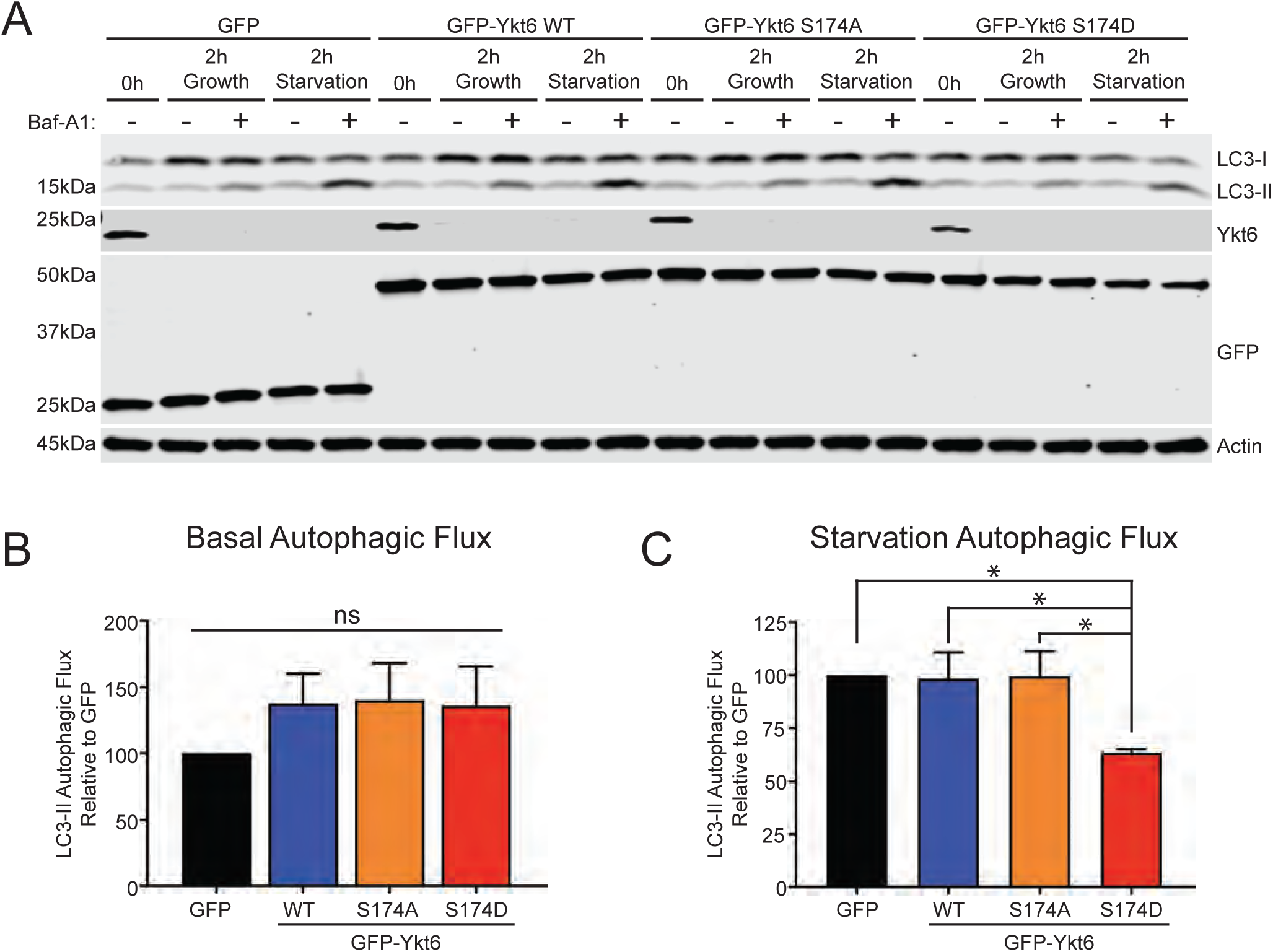
Ykt6 effects on autophagosome/lysosome fusion depend on the phosphorylation at the evolutionarily conserved site in the SNARE domain in the absence of endogenous Ykt6. (A) Representative western blot for LC3-II from a stably transfected PC12 cell line with a doxycycline inducible shRNA targeting endogenous rat Ykt6 and infected with lentiviruses carrying GFP-tagged wild-type (WT) human Ykt6 and phosphomutants. Cells were treated for 2 hours (2h) with 200nM Bafilomycin A_1_ (Baf-A_1_) in growing conditions (fresh 10% FBS and 4.5% glucose growth medium) or in starvation conditions (Torin-1 250nM, 1X HBSS supplemented with 10mM HEPES). Actin serves as a loading control. **(B-C)** Autophagic flux (defined as [LC3-II Baf-A_1_] – [LC3-II]) from western blot in (A) of **(B)** growing and **(C)** starvation conditions. N=5 *p<0.05 **p<0.01 ****p<0.0001 One-way ANOVA, uncorrected Fisher’s LSD test.

## DISCUSSION

While most SNAREs are permanently anchored to membranes via their transmembrane domain, Ykt6 membrane localization is regulated by reversible lipidation at the C-terminus (Fukasawa et al., 2004). Ykt6 activation has been proposed to be regulated via a conformational change from a closed cytosolic form to an open membrane-bound form, although specific mechanisms and/or factors driving these transitions were not known. Through biochemical, structural, genetic and pharmacologic approaches, we found that a critical determinant for Ykt6 conformational switch and membrane association is driven by phosphorylation. We showed that phosphorylation at an evolutionarily conserved site within Ykt6 SNARE domain can regulate a conformational change from a closed cytosolic form to a more open membrane-bound form. Furthermore, our data suggests that the negative charge introduced by phosphorylation creates additional electrostatic potential of the protein which drives the separation of the hydrophobic Longin and SNARE domains (Figure 9).

**Figure 9.**
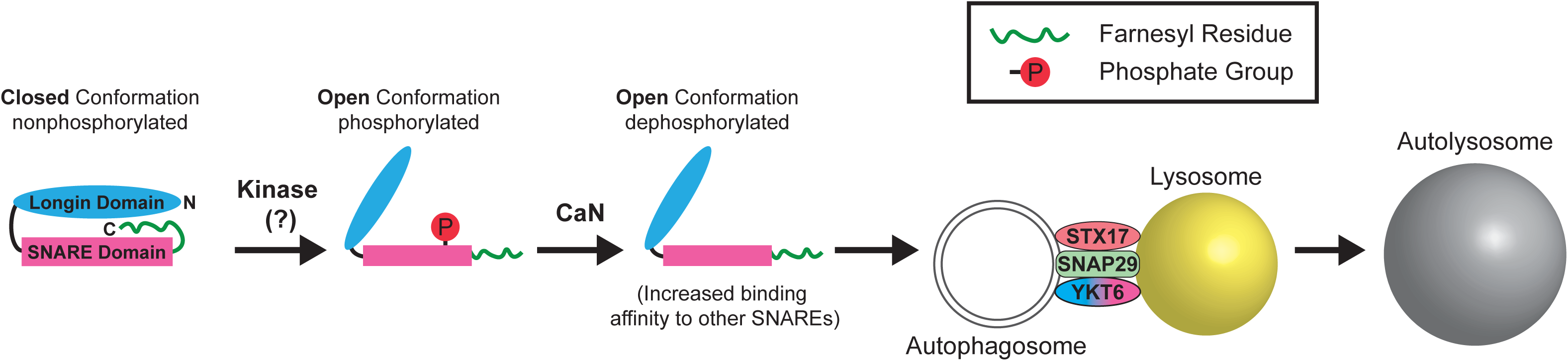
Models depicting the role of phosphorylation in Ykt6 SNARE domain and its impact in conformation. Diagram representing the effect of Ykt6 phosphorylation in its conformation and activity during autophagy. Ykt6 exists in a closed conformation in the cytosol and a rise in Ca^2+^, in this case due to starvation, activates an unknown Ca^2+^-dependent kinase which phosphorylates Ykt6 in the evolutionary-conserved calcineurin-sensitive site promoting its open conformation. Subsequently, an increase in calcineurin (CaN) activity, also due to increased cytosolic Ca^2+^, dephosphorylates Ykt6 promoting interactions with SNAREs involved in autophagy (such as SNAP29 and STX17) thereby promoting autophagosome/lysosome fusion.

We found that this conformational switch impacts the Ykt6 interactome. We showed that Ykt6 conformational change induced by phosphorylation is an important regulatory step that mediates binding of critical members of the autophagosome/lysosome fusion machinery to complete autophagy. In Drosophila cells, Ykt6 was recently shown to participate in autophagosome/lysosome fusion in a complex with SNAP29 and Stx17 as a scaffold (Takats et al., 2018). This Ykt6 non-SNARE mediated function was based on the finding that a Ykt6 mutant with a Q residue in the zero layer instead of the native R could still perform autophagy (Takats et al., 2018). In mammalian cells, we found that phosphorylation within Ykt6 SNARE domain governs the interaction with Stx17 and SNAP29 and is a key determinant for Ykt6 function in autophagy suggesting that in mammalian cells this is a SNARE mediated interaction. Another group also recently shown that Ykt6 participates in autophagosome/lysosome fusion by forming a complex with SNAP29 and Stx7 in mammalian cells (Matsui et al., 2018). The presence of two autophagosome/lysosome complexes is not necessary conflicting but rather can constitute an additional layer of regulation to assure autophagy completion is efficiently achieved. Our data would explain why Stx17 and Ykt6 are not simply redundant but rather have an additive effect as another group reported (Matsui et al., 2018). Alternatively, these two complexes could be involved in mediating fusion of different populations of autophagosomes/lysosomes. For example, one complex could be involved in fusion of autophagosomes with amphisomes and another could be involved in the homotypic fusion of autophagosomes.

We demonstrated that the conserved phosphorylation within Ykt6 SNARE domain is regulated by the highly conserved Ca^2+^-dependent phosphatase calcineurin. Because calcineurin senses Ca^2+^ concentrations and transduces that information into cellular responses, it suggests that the phosphorylation site we found is tightly coordinated to cellular demands. One such cellular demand in which cytosolic Ca^2+^ is elevated is autophagy (Decuypere, Bultynck, & Parys, 2011; Sun, Xu, Wang, & Zhang, 2016). While it is known that starvation triggers a Ca^2+^-calcineurin cascade to transcriptionally activate autophagy (Medina & Ballabio, 2015; Medina et al., 2015), our data suggests that calcineurin can regulate autophagy through post-translational regulation of Ykt6 activity. Defects in Ca^2+^ homeostasis and autophagy have been implicated in Parkinson’s Disease, the second most common neurodegenerative disease (Cerri & Blandini, 2018; Surmeier, Obeso, & Halliday, 2017). Defects in Ykt6 function have also been linked to the pathobiology of *α*-syn (Cooper et al., 2006; Thayanidhi et al., 2010), a small lipid binding protein whose misfolding is the hallmark of both familial and sporadic Parkinson’s Disease (Polymeropoulos et al., 1997; Simon-Sanchez et al., 2009; Singleton et al., 2003; Spillantini et al., 1997). How Ykt6 affects neuronal health in these models remains poorly understood. Therefore, our studies on Ykt6 regulation can shed light into key aspects of Parkinson’s Disease pathology and potentially provide new therapeutic routes for the disease.

Our findings suggest a model in which an elevation of intracellular Ca^2+^ triggered by autophagy leads to phosphorylation of Ykt6 in the SNARE domain by a yet unknown Ca^2+^-dependent kinase promoting Ykt6 open conformation (Figures 3A,B and Figure 9). We noticed that the additional calcineurin-dependent phosphorylation sites retrieved from the human mass spectrometry analysis (S172 and T179) are also located in the two C-terminal helices of the Ykt6 SNARE domain, *α*F and *α*G, which are more exposed to solvent compared to S174 (Supplemental Figure 9A). These phosphorylation sites, while they are not conserved in yeast, are highly conserved in mammals (Figure 1B). Similar to S174, placement of phosphate at these sites did not cause an obvious clash of the atoms (Supplemental Figure 9B). However, the electrostatic potential became even more negative compared to the single phosphate at S174 (Supplemental Figure 9C). The evolutionarily conserved S174 is less exposed to solvent relative to the other sites, but it is in closer proximity to the negatively charged patch of residues within the loop. This may indicate that S172 and T179 are triggers of the open conformation, while S174 may play a key role in maintaining the open conformation. Phosphorylation of the less evolutionarily conserved sites could be a first step to open Ykt6 followed by phosphorylation of the evolutionarily conserved site S174 to stabilize the open conformation and shift it into a membrane (Figure 9). In a second step, activation of calcineurin would dephosphorylate Ykt6 at the evolutionarily conserved site S174 in the SNARE domain, thereby enabling Ykt6 interaction with other SNAREs such as SNAP29 and Stx17 facilitating autophagosome/lysosome fusion (Figure 9). This would explain why the phospho-ablative S174A had more interactions with SNARE proteins reflecting the fact that Ykt6 can still be somewhat opened and subsequently readily incorporated into the hydrophobic core for SNARE complex formation due to the complete lack of phosphate at S174.

Moreover, this two-step model would help explain why S174 has the highest degree of conservation across phyla. It is interesting to note that the evolutionarily conserved S174 phosphorylation site we found in the SNARE domain is not only restricted to Ykt6, but also evolutionarily conserved across other R-SNARE proteins (Malmersjo et al., 2016). This suggests that the conformational change caused by phosphorylation within the SNARE domain might be a general regulatory mechanism of other SNAREs and prompts further investigation. While we have focused our study on the role of Ykt6 phosphorylation in autophagy, it remains to be determined whether phosphorylation at the evolutionarily conserved site can affect other essential and novel Ykt6 phosphorylation-dependent interactions highlighted by our mass spectrometry data.

## METHODS

### Cells line and manipulations

All cell lines were cultured at 37°C and 5% CO^2^. HEK293T (human embryonic kidney cells) and HeLa (human cervical cancer cells) cells were cultured in Dulbecco’s Modified Eagle Medium with 4.5% glucose (DMEM; Corning), 10% Fetal Bovine Serum (FBS; Denville) and 1X penicillin and streptomycin (Gibco). PC4 cells (Gordon et al., 2017) were cultured in DMEM, 10% FBS and puromycin (1 µg/mL, Sigma Aldrich). CMV-GFP-LC3 stable HeLa cell line was cultured as described above with G418 (0.1 mg/ml, Roche). PC12 (rat pheochromocytoma) cell line was stably infected with a lentivirus carrying a doxycycline inducible rat Ykt6 shRNA (targeting sequence: CAGTCGAAAGCCTTCTATA, catalogue number V3SR11254-241803912 from Dharmacon) and cultured in DMEM, 1% FBS, 10% Nu-serum (Corning), 1X penicillin and streptomycin, and puromycin (1 µg/mL, Sigma Aldrich).

Transfections were performed using either Jetprime (VWR; 89129-924) or Lipofectimine3000 (Thermofisher; L3000015) according to the manufacture’s protocol. Cells were seeded at 55K cells/well in a 24-well plate, and 300K cells/well in a 6-well plate. Autophagy was induced by treating the cells for 2 hours in 250nM Torin-1 (R&D Systems, 4247/10) in 1X Hanks’ Balanced salt solution, HBSS, with Calcium and Magnesium (Corning, 21-020-CV) supplemented with 10 mM HEPES buffer (Corning, 25-060-CI) at 37°C with 5% CO^2^. To block lysosomal degradation cells were treated with 200 nM Bafilomycin A_1_ (Baf-A_1_, Cayman Chemical, 11038) for 2 hours. For all calcineurin related experiments, cells were treated for 30 minutes with 1 µM of Ca^2+^ ionophore ionomycin (Sigma, 407952-1MG) and/or 1 µM of calcineurin-specific inhibitor Tacrolimus (Ontario Chemicals Inc., 104987-11-3).

### Virus Packaging

HEK293T cells were transiently transfected as described above with three constructs: pMD and pax2 (both required for viral packaging), as well as the desired lentiviral construct being packaged. Transfected cells were incubated at 37°C and 5% CO^2^ for 48 hours and supernatants were collected and spin-cleared from cell debris at 500g for 10 min.

### Spin Infection

PC12 cell media was exchanged for packaged virus-containing media supplemented with 0.8µg/mL polybrene. PC12 cells were subsequently spun at 2250RPM for 2 hours at room temperature. After spin, virus-containing media was discarded and replaced with PC12 full-growth media. PC12 cells were assayed 48 hours post infection.

### PC12 rat Ykt6 shRNA Stable Cell Line

PC12 cells were infected as described above (spin Infection) and 48 hours post-infection treated with puromycin (1 µg/mL) for selection of the ShRNA.

### Ykt6 Knockdown

Ykt6 shRNA stable PC12 cell line was infected with GFP-tagged human Ykt6 and mutants as described in Virus Packaging and Spin Infection. After 2-hour spin infection, virus-containing media was discarded and replaced with PC12 full-growth media supplemented with 10ng/mL doxycycline for 48 hours to allow the induction of the shRNA.

### PC4 Secretory assay

PC4 cells (Gordon et al., 2017) were incubated with 0.5 µM D/D Solubilizer (Takara; 635054) in a time course-dependent manner. D/D Solubilizer was deactivated by incubating the cells for 10 minutes on ice and subsequently trypsinized in 0.25% EDTA (Corning) on ice for an additional 30 minutes. All supernatants were collected and analyzed by fluorescence activated cell sorting (FACS) for both, mcherry (Ykt6 transfected) and GFP (secretory reporter) positive cells.

### Yeast maintenance and manipulations

We utilized two types of yeast strains: W303 (for the overexpression experiments) and BY/S288c Ykt6 Temperature sensitive strain (Ts) (Kofoed et al., 2015). To transform the Ykt6 constructs into both strains, yeast cells were grown overnight to saturation at 30°C in a shaking incubator. The following day, the culture is diluted to 0.1 OD_600_. When cultures reach 0.4-0.6 OD_600_, they were centrifuged at 4,000g and washed with autoclaved deionized water. The final pellet was resuspended in 0.1 M lithium acetate (LiAc) and incubated at 30°C for 15 minutes, followed by the addition of the plasmid DNA, salmon sperm DNA mixture along with a 50% polyethylene glycol (PEG) solution (4 mL 50% PEG; 0.5 mL 1 M LiAc; 0.5 mL water). Cells+DNA were incubated at 30°C for 30 minutes, and 42°C for 22 minutes, then centrifuged and washed of any remaining PEG solution. Transformants were spread onto plates with synthetic dropout (SD) medium containing glucose and lacking relevant amino acids for selection. Plates were incubated at 30°C for 3 days. For <u>spotting assays</u>, yeast cell cultures were grown overnight at 30°C in SD medium lacking the relevant amino acids and containing glucose. Cell concentrations (OD_600_) were adjusted to the concentration 0.05 OD_600_, five-fold serially diluted, and spotted onto SD medium plates containing either glucose (uninduced) or galactose (induced). For the overexpression experiments, plates were incubated at 30°C and 37°C (for the Ts strain) for 2 days (glucose) or 3 days (galactose).

### Plasmids and primers

ER-RFP was purchased from Addgene (62236). Yeast Ykt6 (Dharmacon, YSC4613), was mutated using Q5 Site-Directed Mutagenesis Kit (NEB; E0554S). Yeast Ykt6 was mutated using the following primers: Forward 5’-GGTGGACAAAGCGGAGTCATT-3’ and Reverse 5’-AAATTATCCAACTTTTCACCTCTTTG-3’ sequences to generate the S176>A substitution and Forward 5’-GATGAGTCATTAACGGCAAGTTC-3’ and Reverse 5’-ATCTTTGTCCACCAAATTATC-3’ sequences to generate the S176>DD substitution. Human eGFP-Ykt6 (a kind gift from Dr. Joseph Mazzulli, Northwestern University), was mutated using Q5 Site-Directed Mutagenesis Kit with the following primers: Forward 5’-GGTGTCCAAAGCCGAGGTGCTGG-3’ and Reverse 5’-AAGTCATCTAGCTTCTCACCTCGC-3’ sequences to generate the S174>A substitution and Forward 5’-GGTGTCCAAAGACGAGGTGCTGG-3’ and Reverse 5’-AAGTCATCTAGCTTCTCAC-3’ sequences to generate the S174>D substitution. mCherry-Ykt6 constructs were made by subcloning Ykt6 mutants from eGFP-Ykt6 plasmids using KpnI and MfeI into the mCherry-C1 (a kind gift of Dr. Gregory Smith, Northwestern University). Split Venus constructs (a kind gift from Dr. Michael Schwake, Northwestern University). To generate the Longin domain fraction, the following primers were used: Forward 5’-AACCGGTATGAAGCTGTACAGCCTCAGC-3’ and Reverse 5’-TTTTCTCGAGTCAAGCTTCTCGTGGGTTCTG-3’. The PCR product for the Longing domain was gel purified, cut with AgeI and XhoI restriction enzymes, and ligated into the N-terminal Venus construct cut with the same restriction enzymes. To generate the SNARE domain constructs, human Ykt6 and its mutants described above were used as templates. The following primers sequences were used: Forward 5’-AACCGGTGATCCCATGACTAAAGTGCAG-3’ and Reverse primers 5’-TTTTCTCGAGTCATGAGTTTTGTTTCCGGGC-3’. The SNARE domain PCR products were cut with AgeI and XhoI restriction enzymes and were ligated into the C-terminal Venus construct cut with the same restriction enzymes. eGFP-Ykt6 viruses were made by subcloning eGFP-tagged human Ykt6 and its mutants using NheI and NotI and ligated into the lentiviral vector pER4. The following primer sequences were used for PCR amplification of Ykt6/restriction sites: Forward 5’-AAAGCTAGCATGGTGAGCAAGGGCGA-3’ and Reverse primer 5’-TGCGGCCGCTCACATGATGGCACAGCAT-3’.

### Antibodies

For immunofluorescence the following primary antibodies were used: GFP (Santa Cruz, sc-9996) and TGN46 (Thermo, pa 1-1069). Secondary antibodies used are the following: Alexa 488 (Invitrogen, a21202 and a21206), Alexa 594 (Invitrogen, a21442) and Alexa 647 (Invitrogen, a281883). For Western Blot the following primary antibodies were used: GFP (Santa Cruz, sc-9996), actin (Abcam, ab6276), Ykt6 (Abcam, ab236583), LC3B (Cell Signaling, #2775), p62 (Sigma, p0067), STX17 (Sigma, hpa001204), SNAP29 (Abcam, ab138500) Na+/K+ ATPase (Sigma, 06-172-I), alpha/beta-Tubulin (Cell Signaling, 2148S), and STX7 (Bethyl, A304-512A). Secondary antibodies used are the following: IRDye680 (Fisher, 925-68070) and IRDye800 (Fisher, 925-32211). Secondary antibodies used are the following: IRDye680 (Fisher, 925-68070) and IRDye800 (Fisher, 925-32211).

### SDS-PAGE/Western blotting and Immunoprecipitations

Transiently transfected HEK293T cells (18-24 hours post transfection) were briefly washed with ice-cold 1X PBS and lysed using a radioimmunoprecipitation (RIPA) assay buffer (50 mM Tris/HCl pH 7.6; 150 mM NaCl; 20 mM KCl; 1.5 mM MgCl2; 1% NP40; 0.1% SDS) for western blot analysis or 1% Triton X-100 lysis buffer (1% Triton X-100, 10% glycerol, 10 mM Tris/Cl pH 7.5; 150 mM NaCl; 0.5 mM EDTA) for immunoprecipitation experiments. In all experiments, lysis buffer was supplemented with the Halt protease and phosphatase inhibiter cocktail (Thermofisher; 78441). Samples were incubated on ice for 30 minutes and pushed through a 27G needle (10 times) to ensure full lysis. Samples were then centrifuged at max RPM (∼20,000 x g) for 20 minutes. Protein concentration was analyzed with the Pierce BCA Protein Assay kit (Thermofisher). For western blot analysis, after the addition of the appropriate amount of the 6X Laemmli Sample Buffer (Bioland scientific LLC, sab03-02) with 5% ß-mercaptoethanol (Sigma) protein samples were boiled and separated on precast 4-20% Criterion TGX Stain-free gels (Bio-Rad) and transferred to a nitrocellulose membrane (Amersham Protran 0.2 µm NC, #10600001). Membranes were blocked with 5% non-fat milk in 1X Tris-buffered saline (TBS) [50mM Tris/Cl pH 7.4, 150mM NaCl] for 1 hour at room temperature. Membranes were subsequently immunoblotted overnight in primary antibody at 4°C, shaking. The following day, membranes were washed three times with 1X TBST (TBS with 0.1% Tween) for 5 minutes and incubated in secondary IRDye antibody for 1 hour shaking at room temperature. Membranes were washed three times with 1X TBST for before imaging using Li-Cor Odyssey® CLx Imaging System. Images were processed using Image Studio Software (LI-COR Biosciences) and densitometries were quantified using Fiji (51). Before final presentation, resulting densitometry values were normalized to the “control”. Autophagic flux is defined as LC3-II (Baf-A1) minus LC3-II (without Baf-A1). GFP-tagged proteins were immunoprecipitated using the GFP-Trap Agarose® beads (Chromotek; gta-20) according to the manufacture’s protocol and eluted with 2x Laemmli Sample Buffer (BioRad; 1610737) with 5% ß-mercaptoethanol for the subsequent western blot analysis. For the mass spectrometry analysis, immunoprecipitated proteins were eluted first with elution buffer I (50 mM Tris/Cl pH 7.5, 2 M Urea, 1 mM DTT) by incubating samples in a thermomixer at 30°C for 30 min and shaking at 400 rpm. Supernatant was collected by spinning at 2500 x g at 4°C for 2 min. Second elution was done with elution buffer II (50 mM Tris/Cl pH 7.5, 2 M Urea, 5 mM iodoacetamide) and repeated once. All collected supernatants were combined and incubated in the thermomixer at 32°C at 400rpm overnight, protected from light. Next day, all samples were submitted for mass spectrometry.

### Partial Proteolysis

HEK293T cells were immunoprecipitated using GFP-Trap beads according to the manufacture’s protocol. After the final spin, the supernatant was discarded and a trypsin (Promega; v5111) solution (25 mM Tris pH 8; 100 mM NaCl; 10% Glycerol; 5 mM MgCl2; Sequence Grade Modified Trypsin) at indicated trypsin concentrations was added to bound-beads and incubated at 25°C for 1 hour. Proteolysis was stopped with a stop solution (500 nM EDTA; 150 mM PMSF; 6X Laemmli Sample Buffer to 400 µL). Samples were boiled and half of the samples was run on the precaste4-20% Criterion TGX Stain-free gels (Bio-Rad) and stain with a Pierce Silver Stain kit (Thermo, 24612) and the second half was analyzed by western blot as described above. Silver stained gels were images using Canon CanoScan 9000F scanner.

### Mass Spectrometry

Relative phosphopeptide quantification by label-free shotgun proteomics was performed in collaboration with Paola Picotti at ETH Zurich as previously described ^33^. Briefly, differentially phosphorylated peptides, we compared control and α-synuclein-expressing yeast cells. We focused on those peptides showing at least a two-fold (log2) change in abundance with a false discovery rate-q value <0.05. Since calcineurin is a phosphatase highly activated in α-syn-expressing yeast cells, we concentrated our analysis on the two-fold hypo-phosphorylated subset (527 of 5,250) to define phosphosites that may be dephosphorylated by calcineurin/FKBP12. Human wild-type (WT) GFP-Ykt6 phospho-peptides were analyzed by isobaric tag for relative and absolute quantification (iTRAQ) mass spectrometry was performed by the MIT Koch core facility. Human Ykt6 WT and phosphomutant interactors were analyzed by the tandem mass spectrometry in Jeffrey Savas’ laboratory at Northwestern University as previously described (Nguyen & Krainc, 2018). Hits were scored as positive based on the following criteria: 1) present in both mass spectrometry analyses, 2) spectral counts had to be at least 2 for Ykt6 WT and/or phosphomutants if the GFP pulldown contained zero spectral counts for a given peptide, and 3) spectral counts had to be at least 10-fold higher than GFP for Ykt6 WT and/or phosphomutants if the GFP pulldown contained greater than zero spectral counts for a given peptide. Sequence alignment was done using Clone manager software (v 9, Sci-ED software) and sequences obtained Uniprot (https://www.uniprot.org/).

### Immunofluorescence and live-cell imaging

HeLa cells were seeded on glass coverslips and fixed with 3% formaldehyde (Fisher 04018-1) 20-24 hours post-transfection for 20 minutes at room temperature. Cells were then permeabilized using permeabilization buffer (10% FBS in 1X PBS + 0.1% fresh saponin) for 1 hour at room temperature. Samples were then incubated in primary antibody diluted in permeabilization buffer overnight at 4°C. Coverslips were then washed three times with 1X PBS for 5 minutes each wash. Secondary antibodies corresponding to the primary antibodies were diluted in permeabilization buffer and added to the samples for 1 hour at room temperature. Samples were washed three times with 1X PBS for 5 minutes each wash and then mounted using a DAPI-staining mounting medium (Vector Laboratories; H-1200-10). Cells were imaged using Leica DMI4000B microscope fitted with Leica TCS SPE laser-scanning confocal system with Solid State lasers (405/488/561/635) and the 40X/1.15 OIL CS ACS APO objective. XY and Z images (10 step, 0.67 micron step) were captured using Leica Application Suite X (LAS X) software. For live-cell imaging, Leica DMI3000B microscope fitted with an QImaging QIClick CCD Camera and Leica HCX PL FLUOTAR L 40X/0.60 CORR PH2 objective was used. Transiently transfected HeLa cells were imaged in VALAB (Vaseline, lanolin, and beeswax) sealed chambers as described previously (Smith, Gross, & Enquist, 2001). Image acquisition was done using Q-capture pro7 software and manual tracking of the equal exposure and digital gain setting between images. All image processing and analysis was done using Fiji (Schindelin et al., 2012). All experiments were done three independent times and analyzed in a randomized and blinded fashion. GFP-LC3-II and split Venus integrated fluorescent intensity was analyzed using CTCF (Corrected Total Cell Fluorescence) = Integrated Density - (Area of selected cell X Mean fluorescence of background readings) formula and normalized to the CTCF value in control cells (fold over control). For the colocalization analysis, Mender’s coefficient was calculated using coloc2 plugin in Fiji. Autophagic flux is defined as LC3-II (Baf-A1) minus LC3-II (without Baf-A1). All values were normalized to the control condition and presented as fold change.

### Structural Analysis

The phosphorylated versions were obtained using Pymol plugin Pytms 52 by modification of the structure of the closed conformation of rat Ykt6 (PDB code 3KYQ) resolved by X-ray resolved diffraction. Model quality was assessed with Molprobity (Williams et al., 2018). According to the evaluation, the generated models have been placed in 100th percentile, arguing that the geometry and contacts of amino acids within the models are comparable to those of the finest current structures in Protein Data Bank. Kinimages visualizing the detailed Molprobity report were generated with KiNG (v2.21). The open conformational state was created by adjusting torsion angles of the backbone in the region 162-164 amino acids using Chimera v1.13 54. Electrostatic potential was computed with APBS plugin of Chimera package. Sequence conservation was analyzed using 196 fungal and 140 animal sequences of Ykt6 proteins retrieved from Tracey database (Pettersen et al., 2004). Sequence logos were created using WebLogo generator (Crooks, Hon, Chandonia, & Brenner, 2004).

### Cell fractionation

HEK293T cells (DMEM, 10 % FBS, 1X P/S) were seeded to ∼80 % confluency a day prior to the experiment in a 10 cm dish. Cells were transfected with empty vector (pEGFP-C1), GFP-tagged-human YKT6 wild type, -S174A and -S174D variants using JetPrime exactly as per manufacturer’s recommendation. Media was replaced after 6 hours and cells were allowed to express the proteins for 24 hours. Subcellular fractionation of transfected cells was carried out exactly as described with no modifications as described in Yu, Z., Huang, Z. and Lung, M. L. (2013). Subcellular Fractionation of Cultured Human Cell Lines. Bio-protocol 3(9): e754.). Fifteen micrograms of the total cell, membrane and cytoplasmic fractions were separated on SDS-PAGE and probed with mouse anti-GFP antibody (Santacruz, 1:2000). Blots were also probed with rabbit anti-Na+/K+ ATPase α-3 antibody (Millipore Sigma, 1:1000) and rabbit anti-tubulin (Cell signaling, 1:10,000) to assess the purity of the membrane and cytoplasmic fractions, respectively. IRDye 800CW goat anti-rabbit and IRDye 680 goat anti-mouse were used as secondary antibodies (Li-COR, 1:10,000). Blots were developed using Odyssey CLx imaging system (LI-COR Biosciences).

## Statistical Analysis

GraphPad Prism 7 Software (http://www.graphpad.com) was used to graph, organize, and perform all statistical analyses. Values are expressed as an average + the standard error mean (SEM). Statistical analysis was determined using the following methods: One-way analysis of variance (ANOVA) with either Tukey’s test or Fisher’s uncorrected LSD test. Two-way ANOVA with Fisher’s uncorrected LSD test. P-values <0.05 were considered significant. A minimum of three biological replicates were used for each experiment. The specific number of biological replicates for each experiment is listed in the Figure Legends.

## ACKNOWLEDGMENTS

For kindly providing critical reagents we would like to thank: Peter Stirling at the University of British Colombia for providing the Ykt6 yeast temperature sensitive strain, Congcong He at Northwestern University for providing the CMV-GFP-LC3 stable HeLa cell line and Andrew Peden at the University of Sheffield for providing the PC4 cell line. Special thanks to Michael Schwake and Robert Vassar for critical reading of the manuscript.

---

**Supplemental Figure 2.**
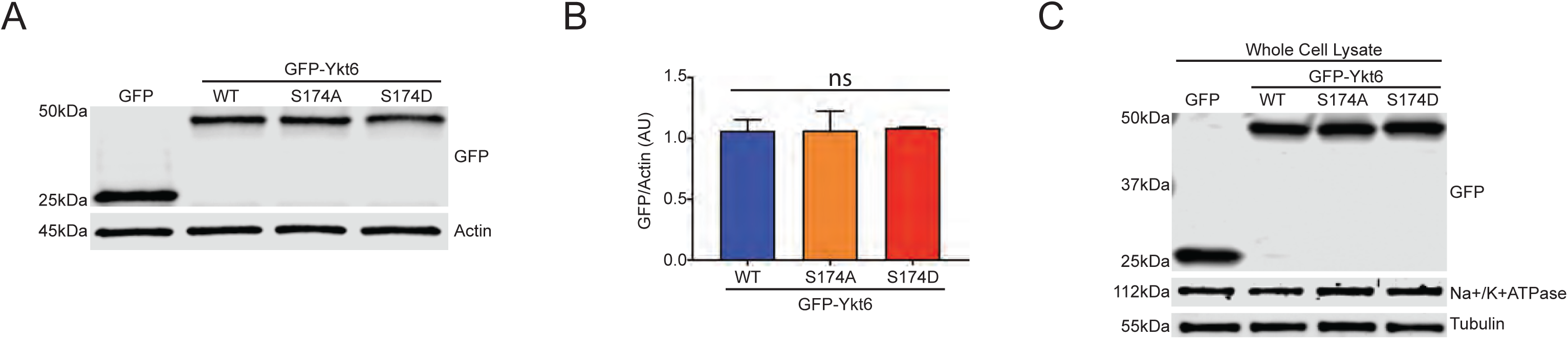
Evolutionarily conserved phosphorylation site within human Ykt6 SNARE domain (S174) is a critical determinant for its intracellular localization. **(A)** Western blot for GFP from transiently transfected HEK293T cells as described in Figure 2A. Actin serves as loading control. **(B)** Densitometry analysis for GFP and actin (loading control) from Western blot in (A) with the indicated Ykt6 constructs. AU=arbitrary units. **(C)** Representative Western Blot for fractionation experiments from Figure 2B of whole cell lysates from HEK293T cells transiently transfected as described in (A).

**Supplemental Figure 4.**
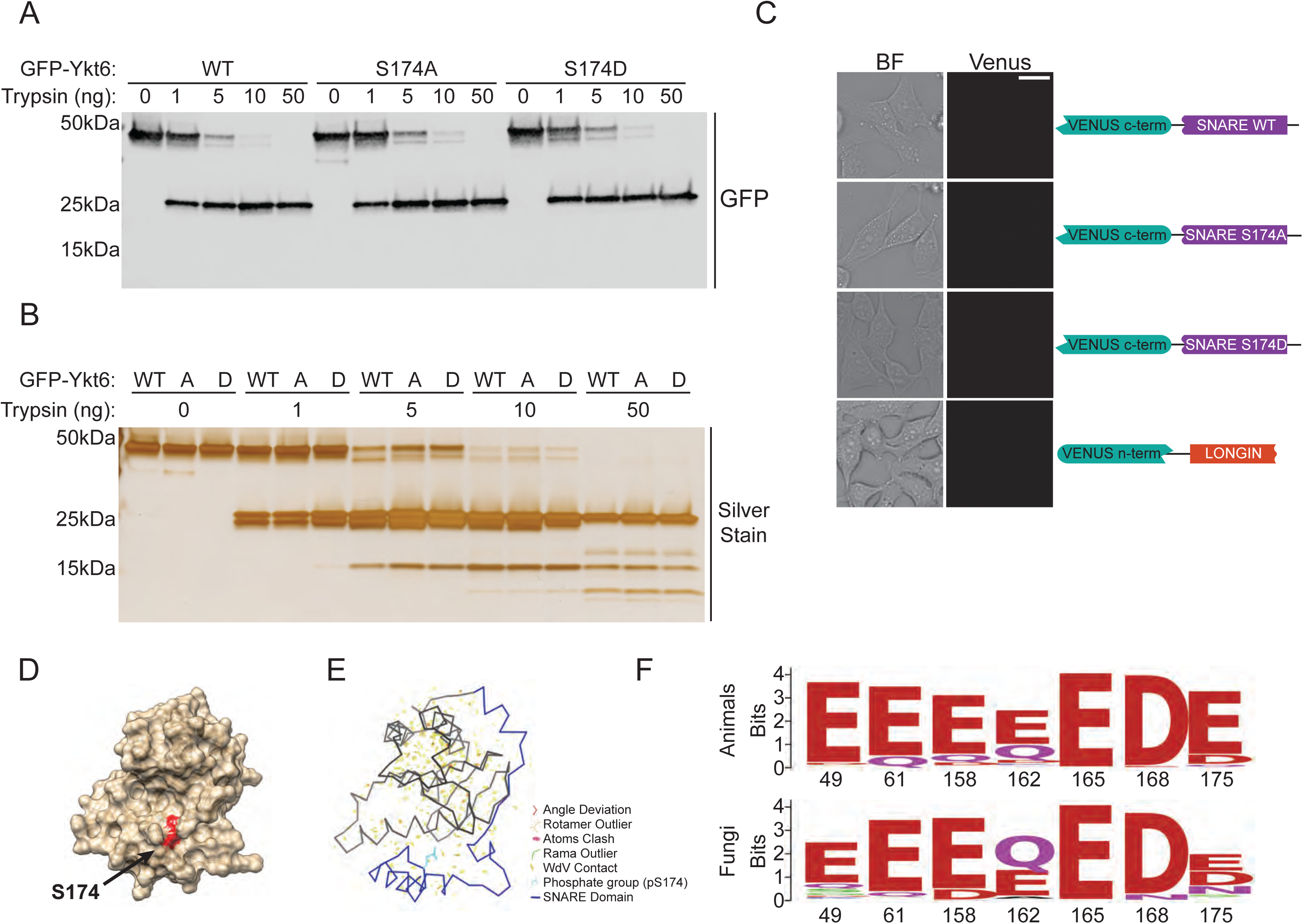
Phosphorylation within the evolutionarily conserved site in Ykt6 SNARE domain regulates its conformation. **(A)** Representative western blot for GFP from GFP-purified wild-type (WT) and phosphomutants of human Ykt6 incubated with the indicated amounts of trypsin for 1 hour at 25°C. **(B)** Representative silver stain gel from GFP-purified wild-type (WT) and phosphomutants of human Ykt6 incubated with the indicated amounts of trypsin for 1 hour at 25°C. **(C)** Representative live-cell phase contrast (BF=Bright Field) and fluorescence images for HeLa cells transiently expressing indicated single split Venus constructs. The Longin domain (LONGIN) and the SNARE domains of either: wild-type (SNARE WT), phosphoablative (SNARE S174A) or phosphomimetic (SNARE S174D) of human Ykt6. Scale bar is 10μm. **(D)** Surface model of Ykt6 closed conformation with the human evolutionary-conserved calcineurin-sensitive phosphorylation site highlighted in red. **(E)** Visualization of atom clashes, poor rotamers, torsion angle outliers using Molprobity and KiNG mimicking the phosphorylation of the human evolutionary-conserved calcineurin-sensitive site in Ykt6 closed conformation with the SNARE domain highlighted in blue. **(F)** Animal and fungal Ykt6 protein sequences were obtained from Tracey database and aligned. The residues corresponding to indicated positions from 3KYQ structure were extracted. Conservation of these residues is shown by sequence logos.

**Supplemental Figure 5.**
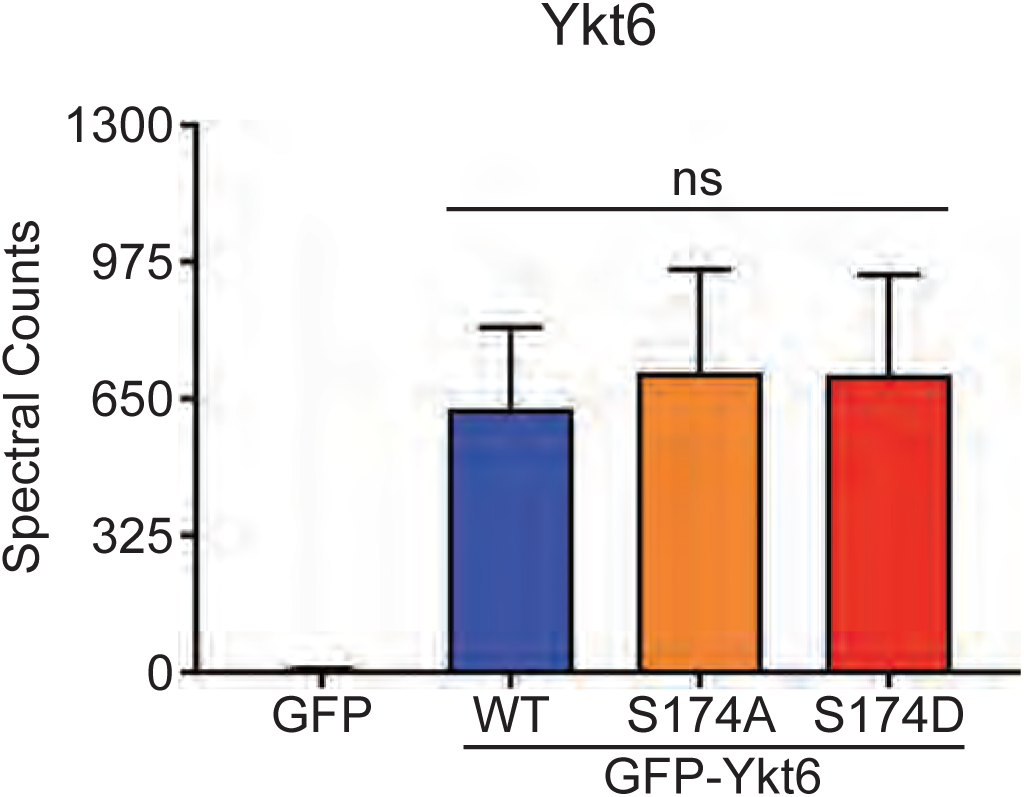
Phosphorylation of the evolutionarily conserved site in Ykt6 SNARE domain S174 affects the specificity of its binding partners. Ykt6 spectral counts from three independent experiments. GFP-Ykt6 wild-type (WT) or GFP-phosphomutants were immunoprecipitated with GFP and subjected to the mass spectrometry.

**Supplemental Figure 7.**
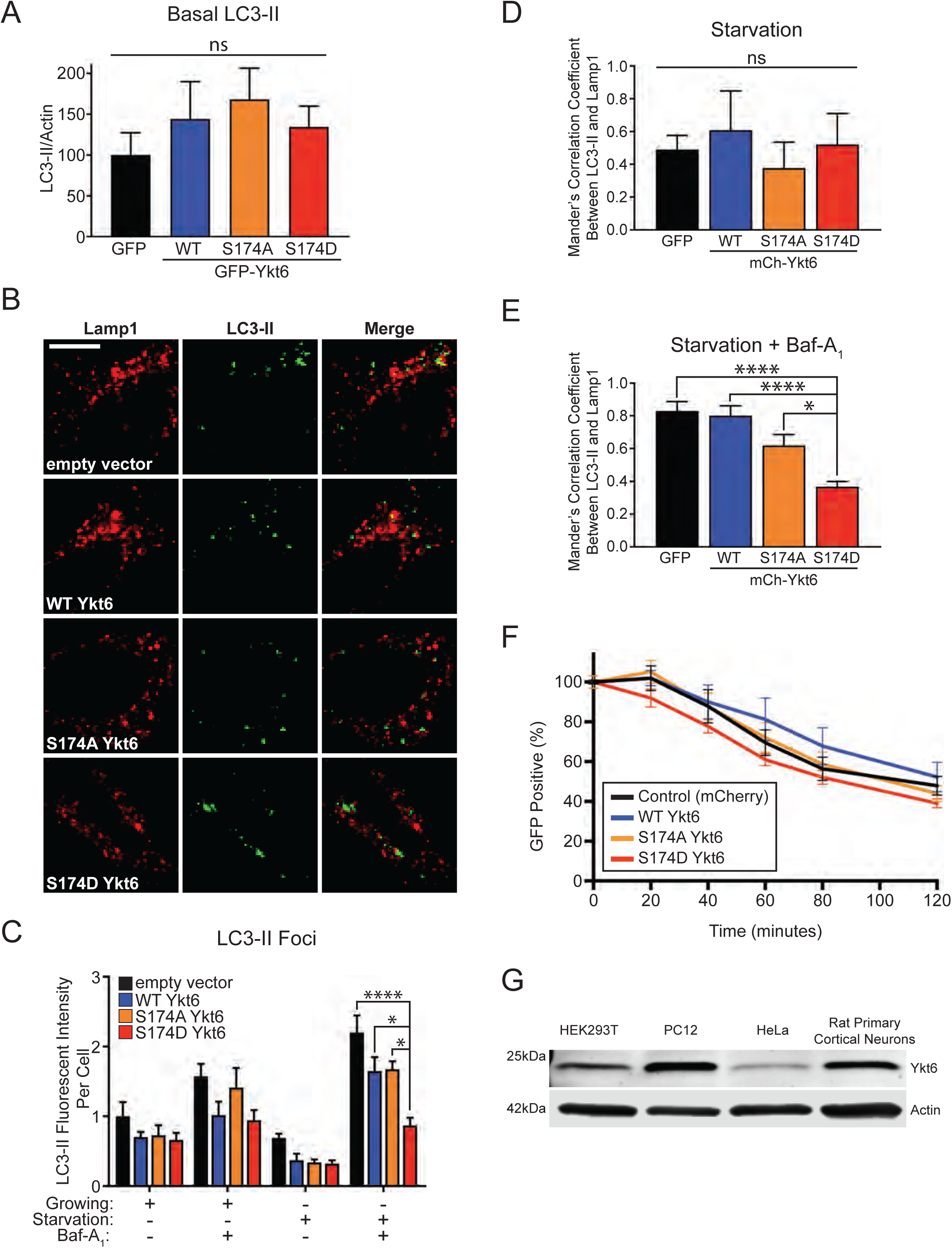
Ykt6 effects on autophagosome/lysosome fusion depend on the phosphorylation at the evolutionarily conserved site in the SNARE domain. **(A)** Densitometry analysis of western blot from Figure 6C of basal LC3-II normalized to Actin. **(B)** Representative immunofluorescence images of overexpressed GFP-LC3 and endogenous Lamp1 colocalization from HeLa cells expressing WT human Ykt6 and phosphomutants, under starvation conditions without Baf-A_1_. **(C)** Normalized fluorescence intensity of GFP-LC3-II foci per cell. **(D-E)** Colocalization analysis based on Mander’s correlation coefficient between Lamp1 and LC3-II in starvation conditions from (B) and in starvation conditions with Baf-A_1_ from Figure 7D. These values were used to calculate autophagic flux graphed in Figure 7F. (A-E) N=3 *p<0.05 **p<0.01 ***p<0.001 ****p<0.0001 One-way ANOVA, Tukey’s test. Scale bar is 5µm. **(F)** Flow cytometry analysis representing the percentage of GFP-FKBP HeLa positive cells transiently expressing either mCherry (control) or N-terminal mCherry fusions of either Wild type (WT), phosphor-ablative (S174A) or phosphor-mimetic (S174D) Ykt6. N=6. **(G)** Representative western blot for endogenous Ykt6 expression in different cell types: HEK293T, PC12, HeLa and rat primary cortical neurons. Actin serves as a loading control.

**Supplemental Figure 9.**
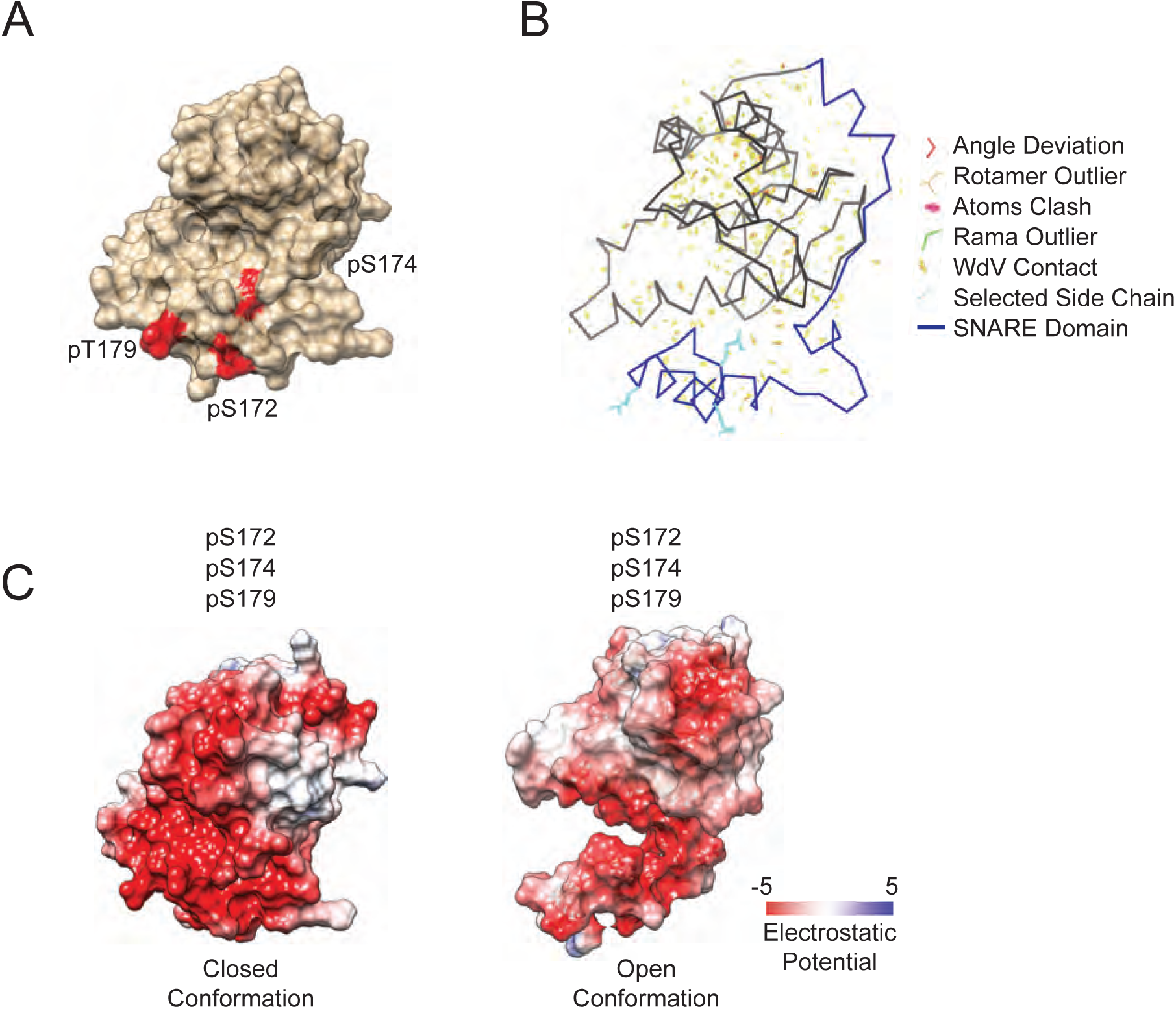
Models depicting the role of phosphorylation in Ykt6 SNARE domain and its impact in conformation. **(A)** Surface model of Ykt6 closed conformation with the human calcineurin-sensitive phosphorylation sites highlighted in red. **(B)** Visualization of atom clashes, poor rotamers, torsion angle outliers using Molprobity and KiNG mimicking the phosphorylation of the human Ykt6 calcineurin-sensitive sites in closed conformation. **(C)** Electrostatic potential computed and projected at the surface representation of both, open and closed conformation, in the absence of phosphorylation (non-phosphorylated), at the evolutionarily conserved site S174 (pS174) and at the additional human calcineurin-dependent sites S172, T179 (pS172 and pT179).

